# CREATE’ing improvements in first-year students’ science efficacy that are independent of instructor rank and experience in a large, multi-section online introductory course

**DOI:** 10.1101/2022.11.14.516496

**Authors:** Jessica Garzke, Blaire J. Steinwand

## Abstract

With a primary objective to engage students in the process of science online, we transformed a long-standing laboratory course for first-year science students into a more accessible, immersive experience of current biological research using a narrow and focused set of primary literature and the CREATE pedagogy. The efficacy of the CREATE approach has been demonstrated in a diversity of higher education settings and courses. It is, however, not yet known if CREATE can be successfully implemented online with a large, diverse team of faculty untrained in the CREATE pedagogy. Here, we present the transformation of a large-enrollment, multi-section, multi-instructor course for first-year students in which instructors follow different biological research questions but work together to reach shared goals and outcomes. We assessed students’: (1) science self-efficacy and (2) epistemological beliefs about science throughout an academic year of instruction fully administered online as a result of ongoing threats posed by COVID-19. Our findings demonstrate that novice CREATE instructors with varying levels of teaching experience and ranks can achieve comparable outcomes and improvements in students’ science efficacy in the virtual classroom as a teaching team. This study extends the use of the CREATE strategy to large, team-taught, multi-section courses and shows its utility in the online teaching and learning environment.

## Introduction

The COVID-19 pandemic forced an unprecedented shift to online learning in higher education (1). The rapid transition to “emergency remote teaching (ERT)” challenged instructors of in-person classes, many of whom were untrained and inexperienced in online teaching practices, across campuses around the world (2). While the remote working and learning environment overwhelmed instructors of all courses traditionally taught on campus, it presented an especially unique challenge to instructors of laboratory courses designed to engage students in the process of science through a “hands-on” curriculum (3).

Laboratory courses give students access to the tools, equipment, and technology involved in the discovery and are, therefore, an integral part of science education (4, 5). Students get the opportunity to participate in the process of science, cultivate problem-solving and critical thinking skills, enhance their ability to understand practical problems and improve their attitude towards learning (6). During the COVID-19 pandemic, science instructors were forced to rethink and creatively redesign laboratory courses to reach these learning outcomes in remote learning environments effectively.

The **C**onsider, **R**ead, **E**lucidate a hypothesis, **A**nalyze and interpret data, **T**hink of the next **E**xperiment (CREATE) strategy focuses on scientific thinking. CREATE uses a novel selection of readings and allows students to engage deeply in activities characteristic of actual science practice (7). In CREATE courses, faculty and students apply evidence-based techniques (e.g. concept mapping, cartooning experiments) in deconstructing and analyzing scientific primary literature. Students learn how to build upon content knowledge and develop the ability to think deeply and critically about the methods and results of scientific studies (8–10). The strategy aims to help students experience the authentic practices of scientists, recognize the limitations of studies and the creativity and open-ended nature of science, and challenge their beliefs about what it means to be a scientist. CREATE allows students to acquire many of the skills traditionally taught in a laboratory setting yet makes participation in the practice of science more affordable, adaptable, and accessible. The student-centered activities that define the CREATE approach, such as concept mapping content, cartooning studies, annotating and transforming data, elucidating hypotheses, and designing experiments, are all conducive to learning online, making the pedagogy a good fit for different learning environments.

The CREATE pedagogy has been introduced and tested at several institutions and studied between two-year and four-year institutions in both introductory and upper-level courses by individual faculty members (7, 8, 11–14). While these studies demonstrate the efficacy of CREATE, they are limited to in-person instruction. At the University of British Columbia (Canada), we redesigned a first-year biology laboratory course for a large-enrollment, multi-section, multi-instructor course for the virtual classroom using CREATE. While different instructors followed different biological research questions, they all used the CREATE approach to reach shared goals and outcomes.

The diversity of teaching experience, research backgrounds, and instructor ranks across the teaching team raised the question of instructor-specific effects on student outcomes. To explore the question of whether an instructor’s degree of teaching experience, research background, and role at the university influenced student outcomes, we grouped instructors of the same rank and title (Research Professor, Research Associate Professor, Associate Professor of Teaching, Assistant Professor of Teaching, full-time contract faculty (Lecturer), course-by-course contract faculty (Sessional Instructor), and Postdoctoral Research Fellow) and compared student outcomes. In this study, we show that novice CREATE instructors with varying levels of teaching experience and ranks (i.e., research faculty, teaching faculty, and postdoctoral research fellows) can achieve comparable outcomes in the affective domain of learning in the virtual classroom when working together.

## Methods

### Context

Participants in this study were undergraduate students at the University of British Columbia (Canada), a large, public, research-intensive university, who enrolled in a term-long “Biology 140: Laboratory Investigations in Life Sciences” course during the winter and spring term of 2020. In total 1,282 students enrolled during both terms (T1: n = 563 students, T2: n = 719 students) were distributed over 52 sections with 25-28 students per section. In total, seven different types of instructors taught a varying number of sections: Research Professors (n = 10), Lecturers (n = 2), Associate Research Professors (n = 2), Assistant Professors of Teaching (n = 1), Associate Professors of Teaching (n = 2), Sessional Instructors (n = 2), and Postdoctoral Research Fellows (n = 3). In each term, before the start of the course, students were invited to participate in a survey voluntarily (with no bearing on class grade) addressing their self-assessed ability to read, analyze, understand, and use a diversity of scientific skills. The same survey was distributed to the students on the last day of the course, allowing us to compare pre- and post-course responses.

### Course Design and Implementation

The CREATE instructional strategy here differed from the original approach (11) designed for upper-level students in which students unpack four papers that follow the same scientific question in their entirety. It also differed from the CREATE Cornerstone approach for first-years (7), in which students analyzed a pair of popular press articles on the same subject. While we used a set of popular press articles to introduce students to a focal topic, we took a deep and focused dive into just one or two figures from two primary scientific articles that followed the same scientific question or real-world biological challenge (Table 1). Literature differed across sections as each instructor focused on a scientific question aligned with their research interest and expertise. This particular aspect of course design engaged our instructors by giving them some autonomy and allowed us to introduce our first-year students to a diversity of research in biology. Although the different topical focus across the course sections varied, all instructors deployed weekly activities associated with the CREATE pedagogy over the course of the term to reach shared learning goals and outcomes despite the different focal topics across sections (Table 1). To establish evidence-based course structure and maintain consistency across sections of the course, we: 1) developed a sandbox site or “shell” on our learning management system that contained all of the shared lesson plans, resources, quizzes, assignments, and activity prompts which we shared with all course instructors for customization before the start of the term, and 2) CREATE’ed an instructor learning community that brought the teaching team together each week for 1 hour to engage in discussions on the course and support one another in the implementation of the pedagogy.

**Table 1.**
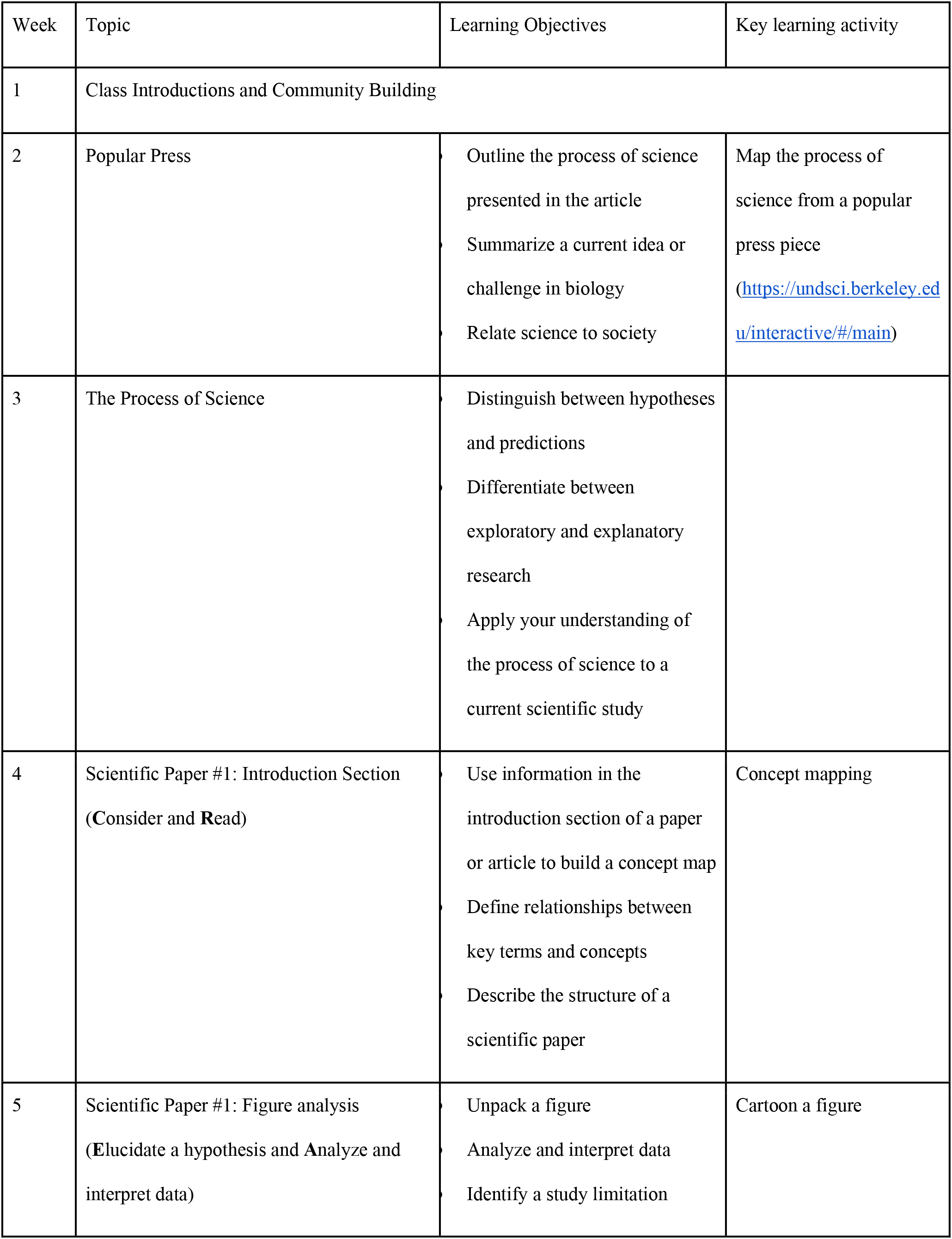
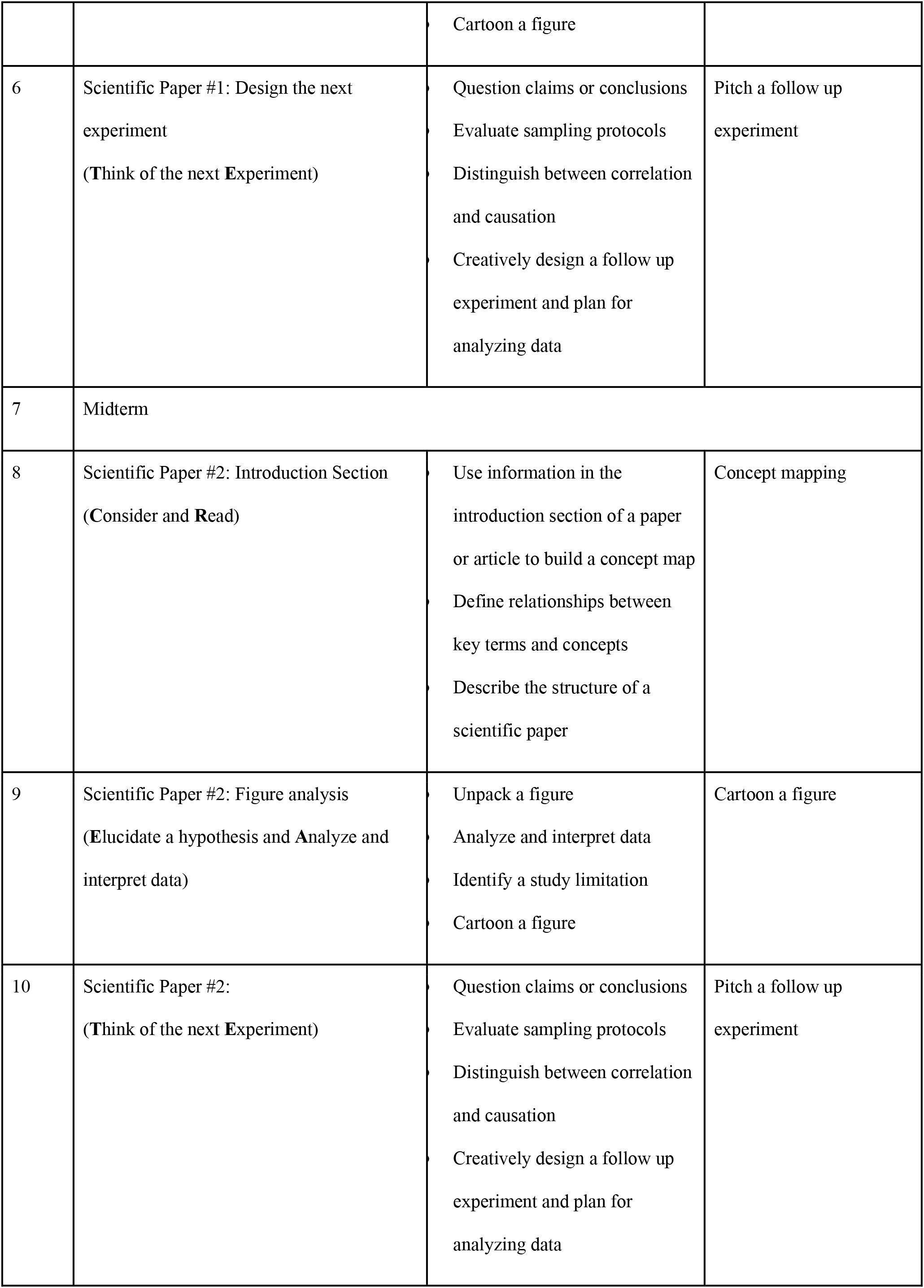
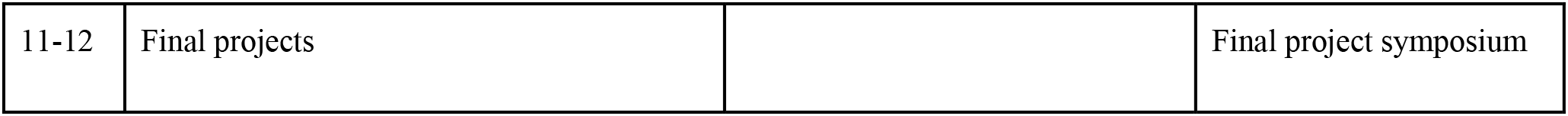
Course schedule of general topics, learning objectives and learning activities

This multi-section course was coordinated by two instructors who planned and facilitated the instructor learning community but needed to gain prior training in CREATE pedagogy themselves. There were 12 instructors on the teaching team in Term 1 and 14 in Term 2. Each week, the instructor learning community (1) reflected on the previous week together, (2) participated in student activities themselves (3) exchanged ideas, resources, and plans for the upcoming week. The instructor learning community allowed all instructors to implement a highly structured, student-centered approach to teaching each week, share teaching practices, and adapt to curriculum and resources as needed. In general, students across all sections of the course were introduced to content and new concepts outside of class sessions. Although early on (typically week 2), some instructors dedicated time to content delivery on their focal topic to help students understand new concepts and the motivation behind the research question, class time was generally structured around CREATE activities that students completed in (online) breakout rooms with members of their fixed groups of 4-5 students.

CREATE activities were implemented online using platforms on which students could work collaboratively, primarily during class time. The platform Miro (miro.com) allowed students to build concept maps of the introduction section of each paper together while students used the media BioRender (biorender.com) and Canva (canva.com) as resources when producing cartoons online (Table 1). All instructors in this study were novices to both the CREATE pedagogy and online teaching.

#### Survey of Student-Rated Abilities, Attitudes, and Beliefs (SAAB)

To assess student outcomes, we administered the SAAB as described in Hoskins et al. (11). The survey consists of 31 statements across seven categories and two individual questions to which students respond on a five-point Likert-style scale (Table 2). An overall score for each of the seven categories reflects pooled responses to the survey statements. The seven individual categories comprise subsets of statements and address specific aspects of scientific thinking and perceptions of science. Surveys were distributed online using Qualtrics in the first (pre-) and final (post-) class sessions and took 15-20 minutes to complete based on the survey platform’s analytics.

**Table 2.** Survey of Student-Rated Abilities, Attitudes, and Beliefs (SAAB). Each question had the same response values available to students: “I strongly disagree” (1), “I disagree” (2), “I am not sure” (3), “I agree” (4), and “I strongly agree” (5), though for the purposes of scoring a number of statements are reverse-coded (i.e., “I strongly agree” would be scored as 1 while “I strongly disagree” would be scored as 5 in such cases). We have indicated reverse-coded statements with an asterisk in this table.

| Question Group | Question |
| --- | --- |
| F1<br>*“Decoding Primary Literature”* | The scientific literature is difficult to understand.* |
|  | When I see scientific journal articles, it looks like a foreign language to me.* |
|  | I am not intimidated by the scientific language in journal articles. |
|  | I am confident in my ability to critically review scientific literature. |
|  | I am comfortable defending my ideas about experiments. |
| F2<br>*“Interpreting Data”* | It is easy for me to transform data, like converting numbers from a table to percentages.* |
|  | If I see data in a table, it is easy for me to understand what it means. |
|  | If I am shown data (graphs, tables, charts), I am confident that I can figure out what it means. |
|  | It is easy for me to relate the results of a single experiment to the big picture. |
| F3<br>*“Active Reading”* | I could make a simple diagram that provides an overview of an entire experiment. |
|  | If I am assigned to read a scientific paper, I typically look at the methods section to understand how the data were collected. |
|  | I do not know how to design a good experiment.* |
|  | The way that you display your data can affect whether or not people believe it. |
| F4<br>*“Visualization”* | When I read scientific material it is easy for me to visualize the experiments that were done. |
|  | If I look at data presented in a paper, I can visualize the method that produced the data. |
|  | When I read a paper, I have a clear sense of what physically went on in a lab/field to produce the results and information I am reading. |
| F5<br>*“Thinking Like A Scientist”* | After I read a scientific paper, I don’t think I could explain it to somebody else.* |
|  | I am confident I could read a scientific paper and explain it to another person. |
|  | I enjoy thinking of additional experiments when I read scientific papers. |
|  | I accept the information about science presented in newspaper articles without challenging it.* |
| F6<br>*“Research in Context”* | Experiments in “model organisms” like the fruit fly have led to important advances in understanding human biology. |
|  | Progress in curing diseases has been made as a result of experiments on lower organisms like worms and flies. |
|  | I understand why experiments have controls. |
| F7<br>*“Knowledge is Certain”* | If two different groups of scientists study the same question, they will come to similar conclusions.* |
|  | The data from a scientific experiment can only be interpreted in one way.* |
|  | Because scientific papers have been critically reviewed before being published, it is unlikely that there will be flaws in scientific papers.* |
|  | Because all scientific papers are reviewed by other scientists before they are published, the information in the papers must be true.* |
|  | Sometimes published papers must be reinterpreted when new data emerge years later. |
|  | Results that do not fit into the established theory are probably wrong.* |

#### Statistical Analysis

SAAB survey response scores were aggregated into the appropriate topic group (Table 2). The median was the average, as the response values do not correspond to a continuous scale. We initially tested, using a two-way repeated ordinal regression model (or cumulative link mixed model), if students of term 1 responded differently than those surveyed during term 2 due to having more experience as a university student in term 2 (R package *ordinal*). A two-way repeated ordinal regression model (or cumulative link mixed model) was used to analyze changes between pre and post-course survey results and between instructor types. Student IDs were coded into random numbers and used as random effects within the ordinal regression model. To identify the best fitting model explaining the observed data, a null model was compared with a model including an interaction term (instructors with pre- and post-survey, Suppl. Table 1). We used the Laplace approximation, logit distribution and equidistant as threshold and identified group differences using the Tukey’s Honest post hoc test comparing the lsmeans (R package *lsmeans* using the sidak approach, Supp Table 2, Suppl. Table 3).

## Results

Initially we tessted if both terms were comparable and if no significant difference occurred; except for question group F4 (“*Visualization*”), in which students in term 1 had significantly higher science self-efficacy compared to students in term 2 (p = 0.285; T1 (mean ± SD): 3.18 ± 0.82; T2 (mean ± SD): 3.03 ± 0.85, Suppl. Table 1), we found no significant differences between terms.

### CREATE shifts students’ self-assessed abilities, attitudes, and beliefs about science when implemented online by a diverse team of novice instructors

We used two “summary” questions from the CREATE survey described in Hoskins (11) to examine students’ overall self-assessed ability to “*read and analyze scientific journal articles*” and “*understanding of the scientific research process*” across all sections of the first-year course. Overall, we observed significant shifts from students feeling less confident in their abilities and understanding pre-term, left-skewed distribution towards feeling more confident in their abilities post-term, right-skewed distribution (Fig. 1A, Table 3). Across all seven question categories of the SAAB survey, we found significant increases in students’ confidence in their science skills, abilities, and epistemological beliefs post-term (Table 3).

**Figure 1.**
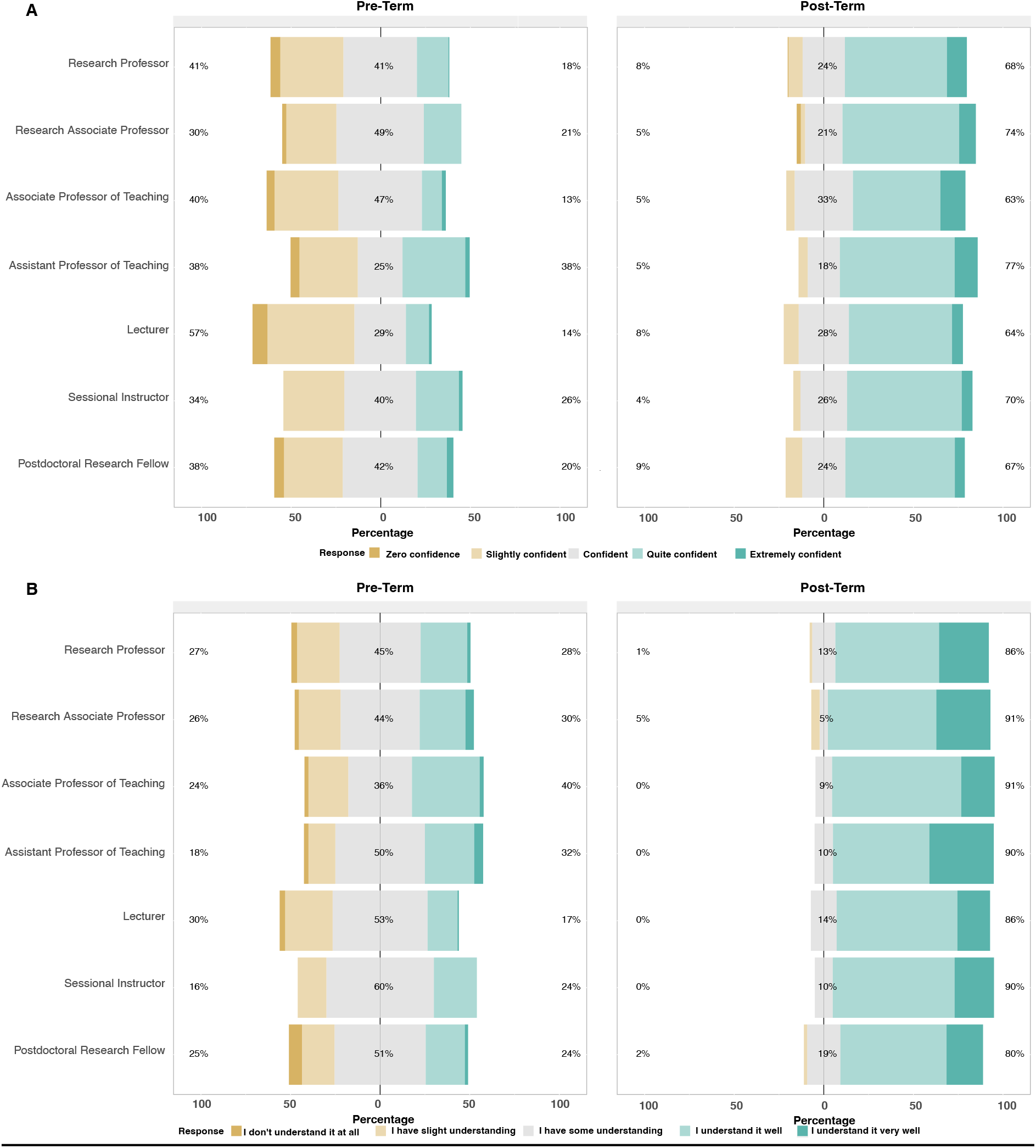
Pre- and post-term SAAB survey results of two summary questions A) “On a scale of 1 to 5, rate your confidence in your ability to read and analyze science journal articles.”, B) “On a scale of 1 to 5, rate your understanding of the way scientific research is done of the scientific research process.”

**Table 3.**
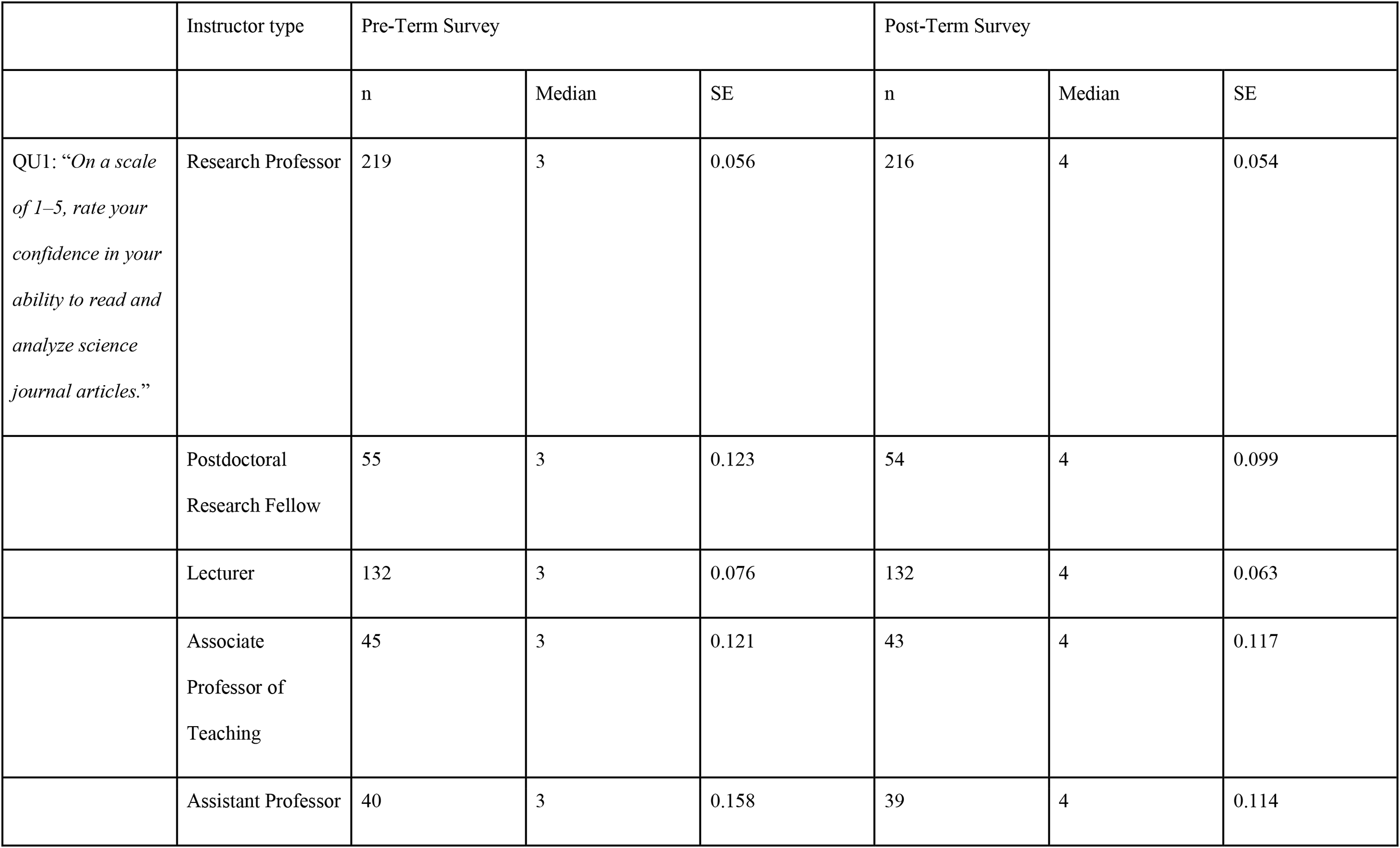
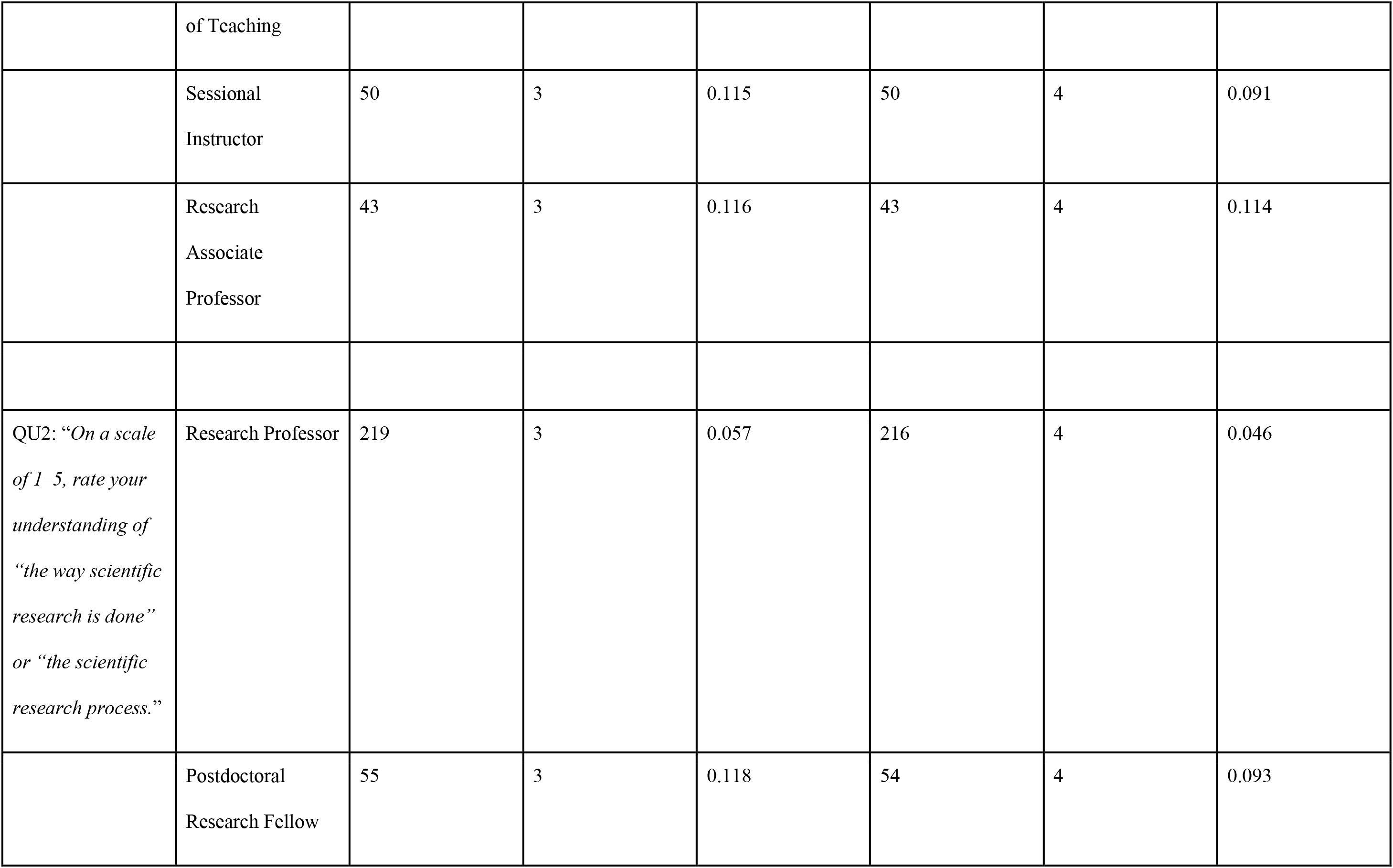
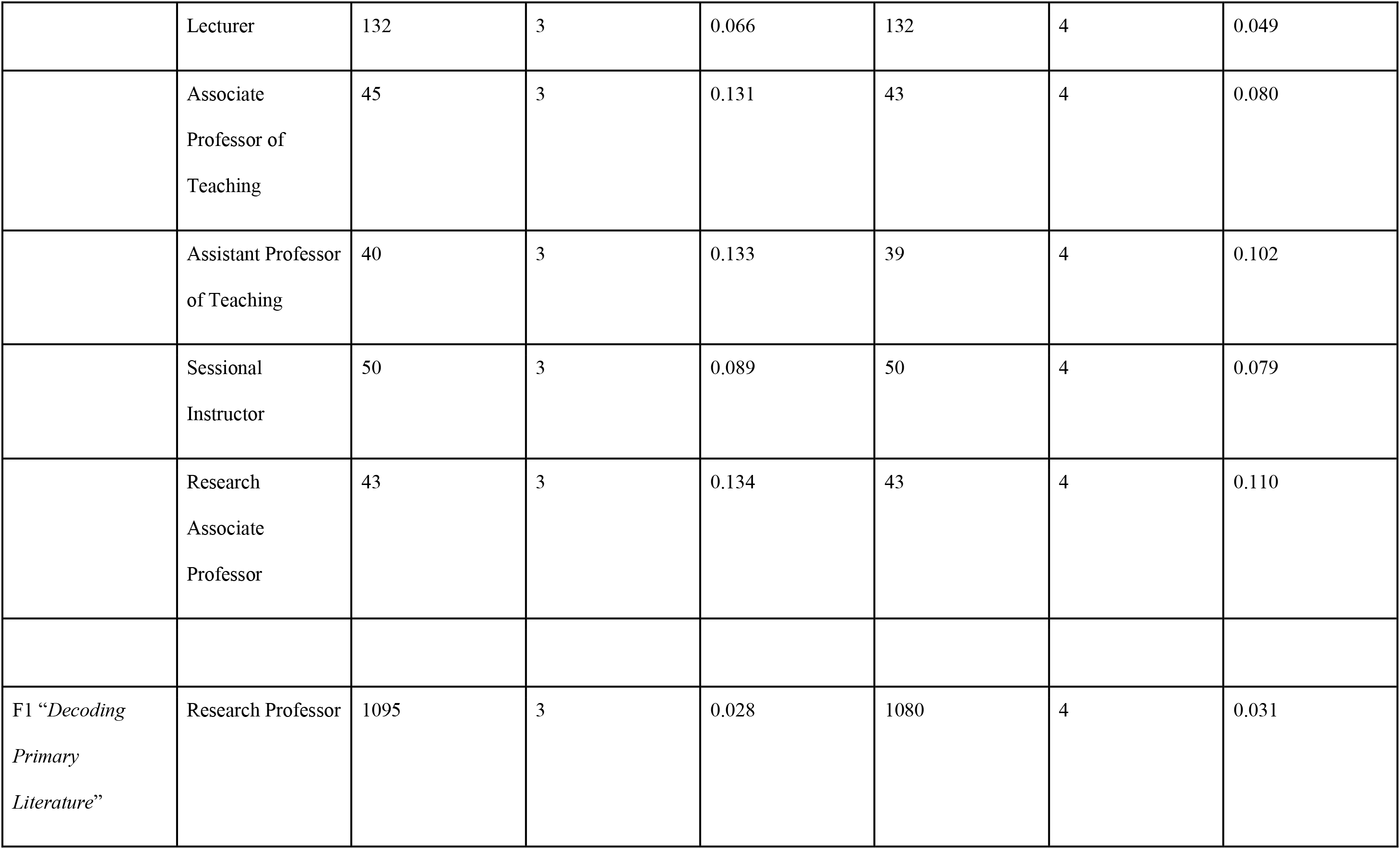
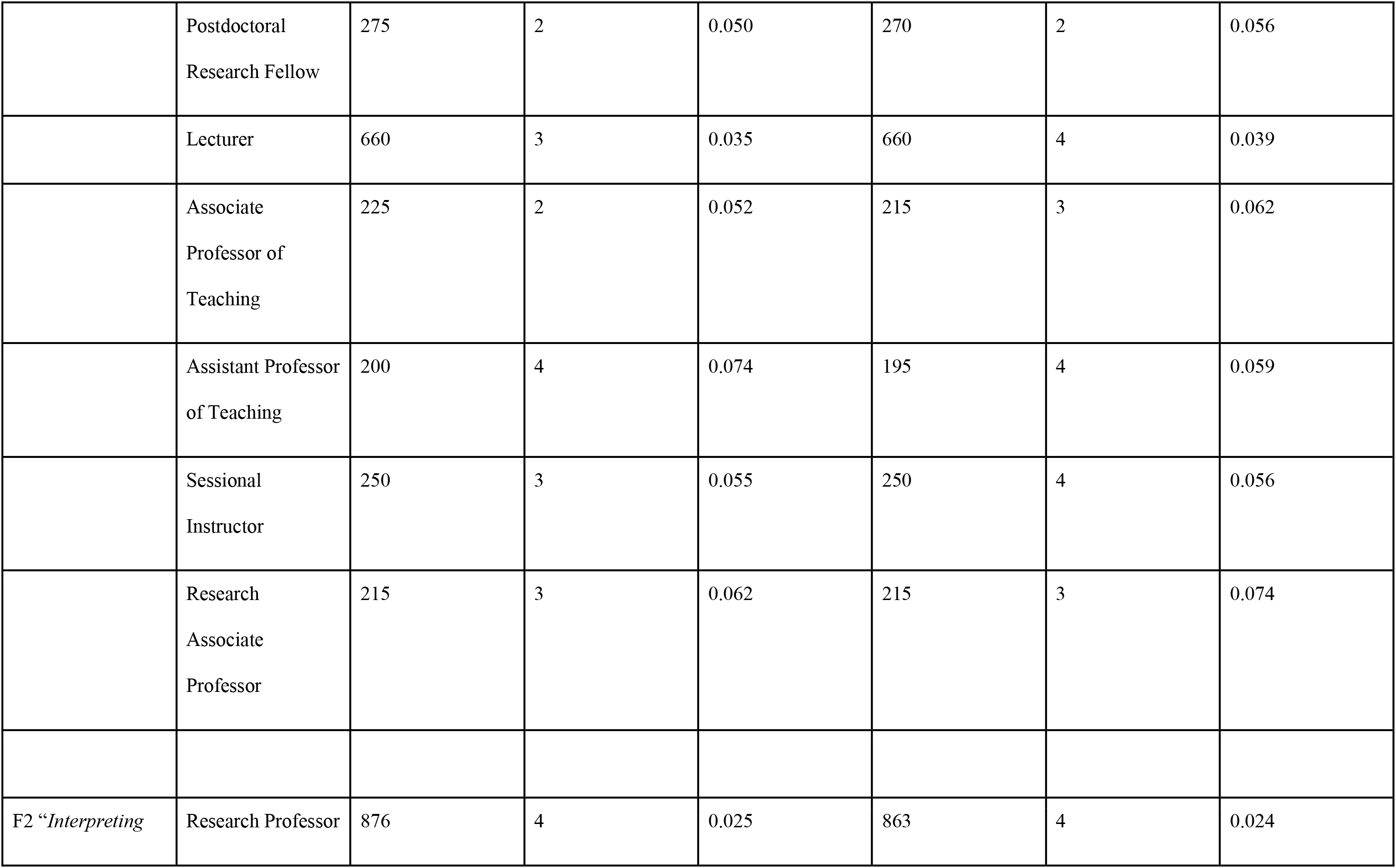
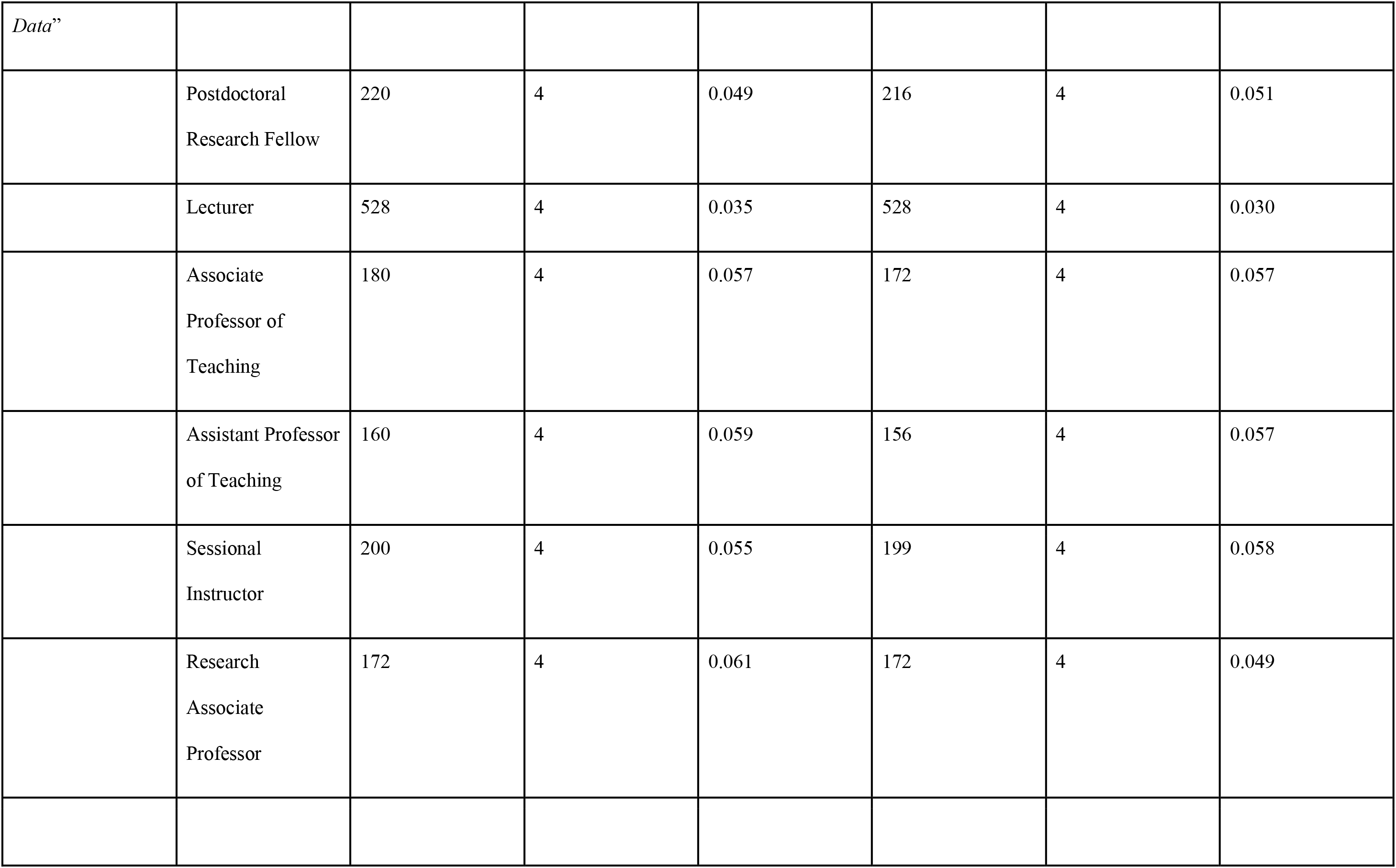
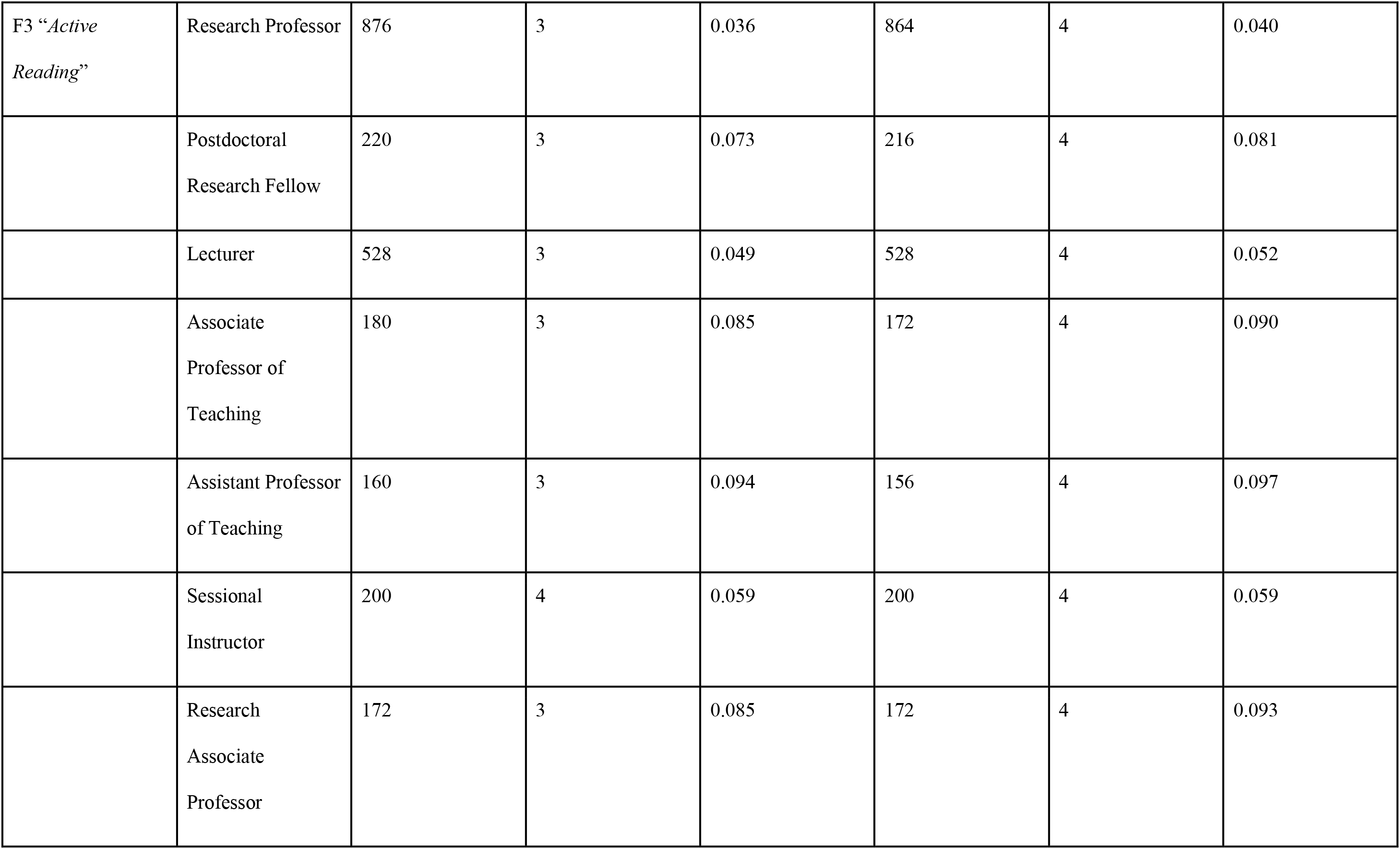
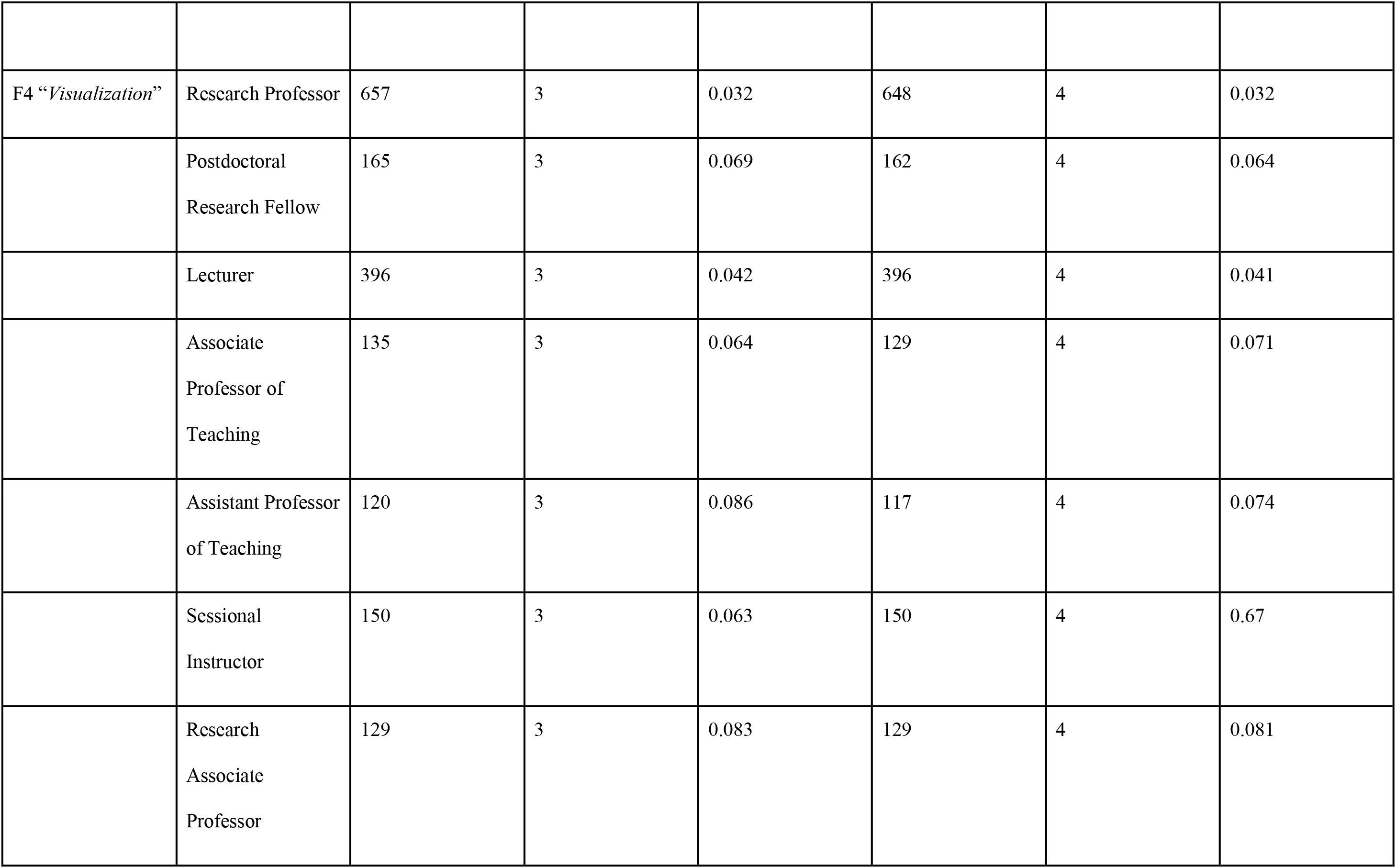
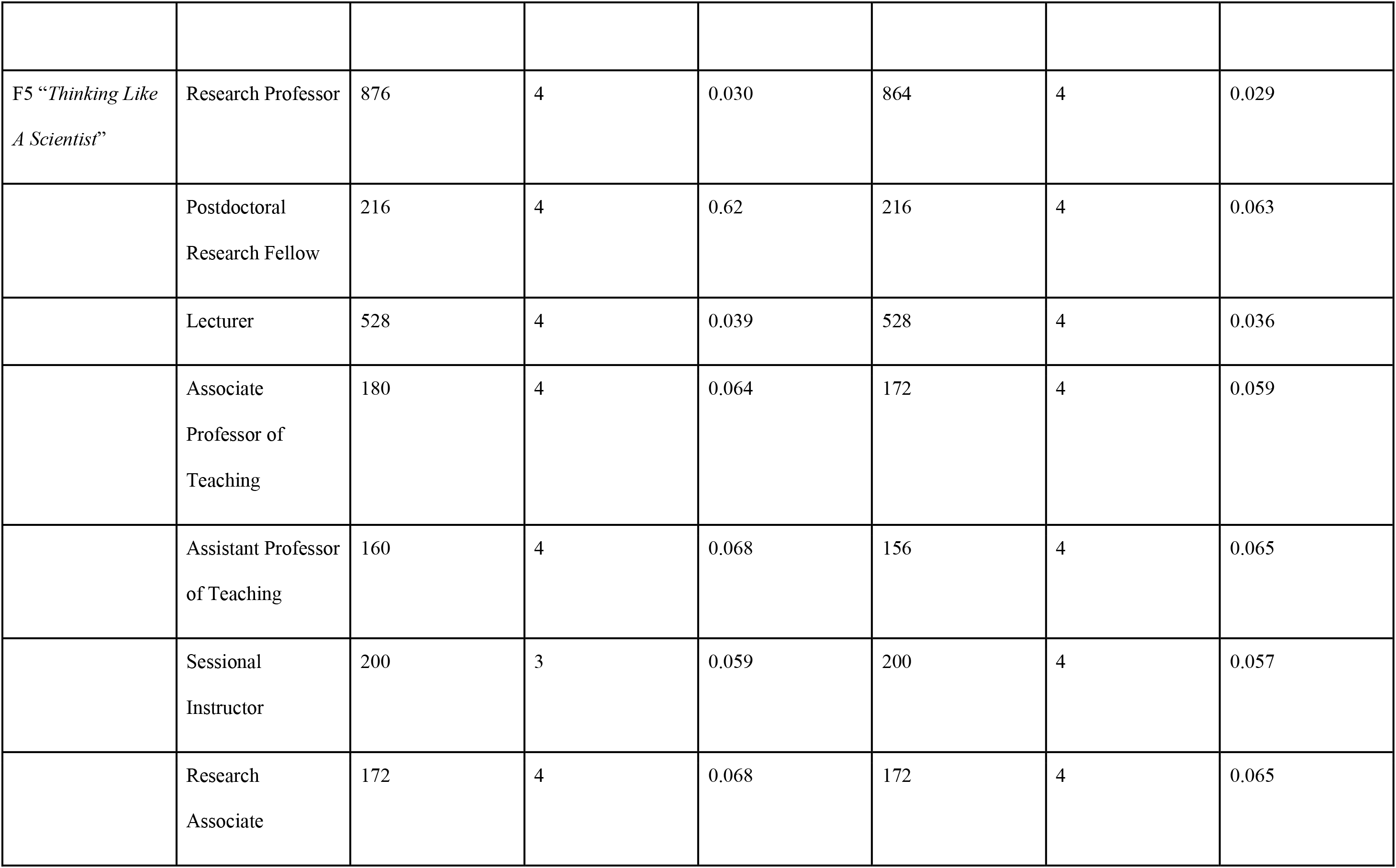
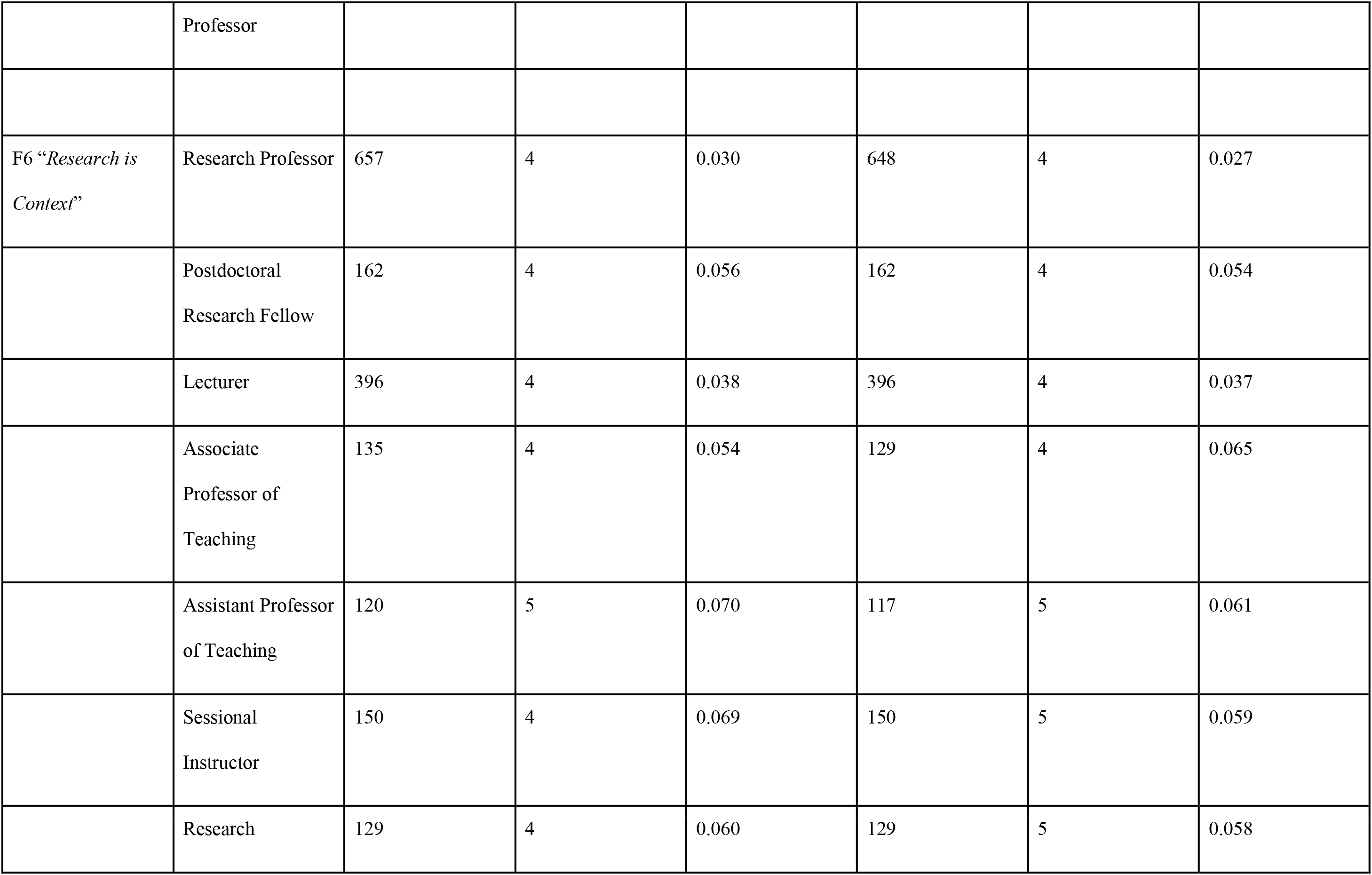
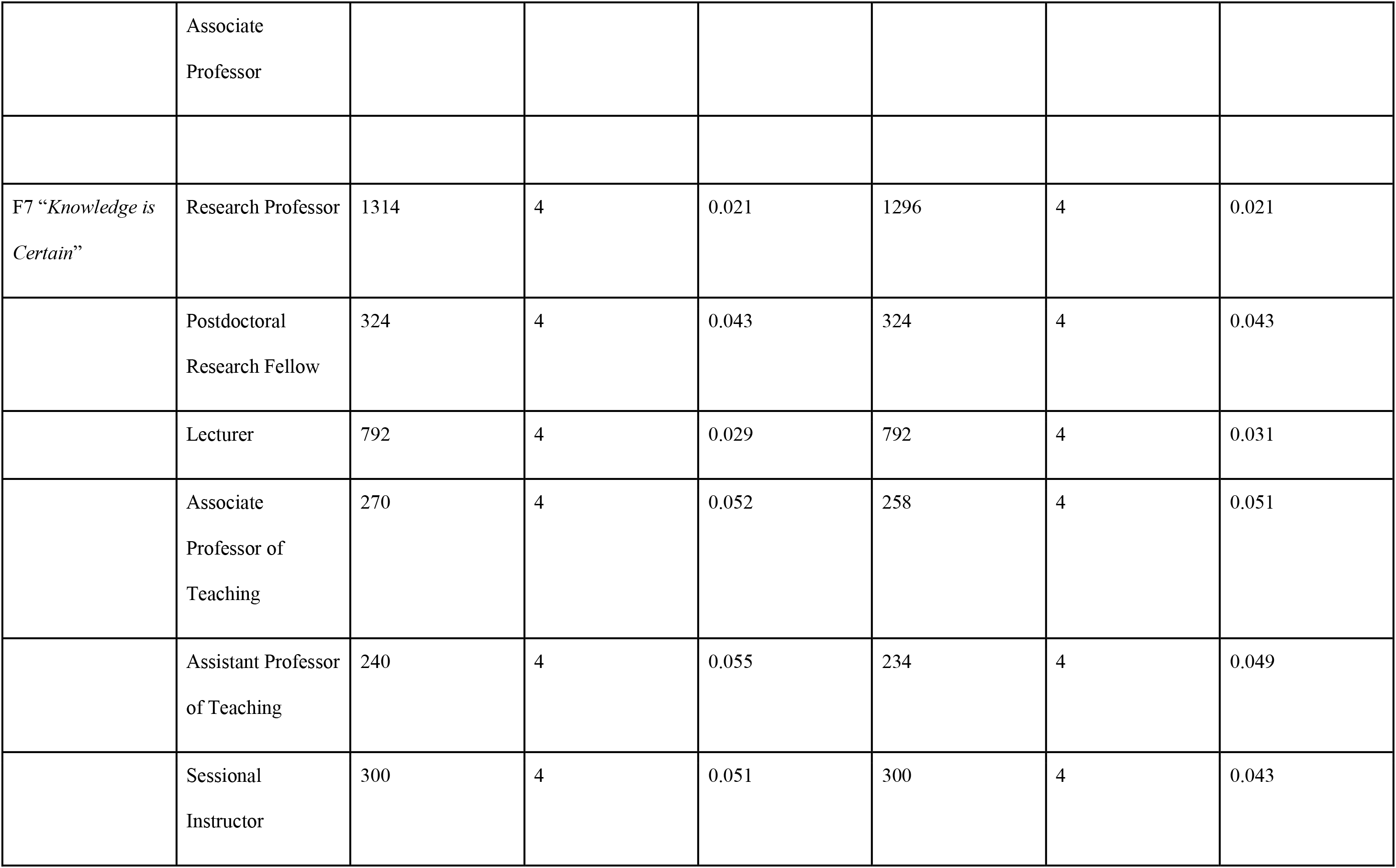
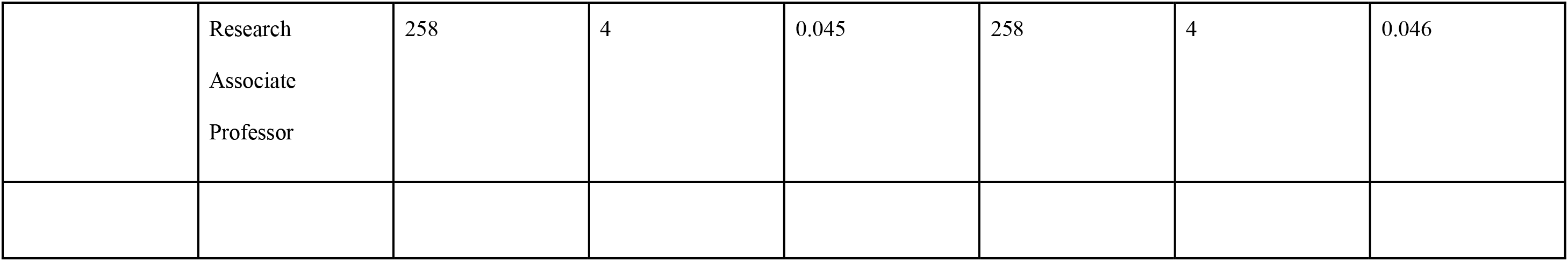
Descriptive analysis of survey results, n = number of students taken the survey, median of all answered given, SE is the standard error.

### SAAB outcomes are not instructor dependent

While students of instructors of all ranks shifted significantly with respect to Question 1 (*“confidence to read and analyze journal articles’’*) and Question 2 (*“rate your understanding of the scientific process”*) (Fig. 1 A and B), we identified some differences between instructor cohorts for individual SAAB categories. We identified two instructor types that experienced slightly smaller, non-significant shifts in student gains and confidence from pre-term to post-term; students of (1) Associate Professors of Teaching and (2) Assistant Professors of Teaching. However, students of both cohorts reported higher confidence in their ability to *decode primary literature* levels pre-term (Fig. 2A). A total of 40% of students taught by Associate Professors of Teaching and 48% of students taught by Assistant Professors of Teaching reported confidence levels of 4 (high) and 5 (very high) coming into the course, whereas 30% – 38% (median 37%) of students in all other instructor types reported high (4) to very high (5) levels of confidence (Fig. 2A, Suppl. Table 2).

**Figure 2.**
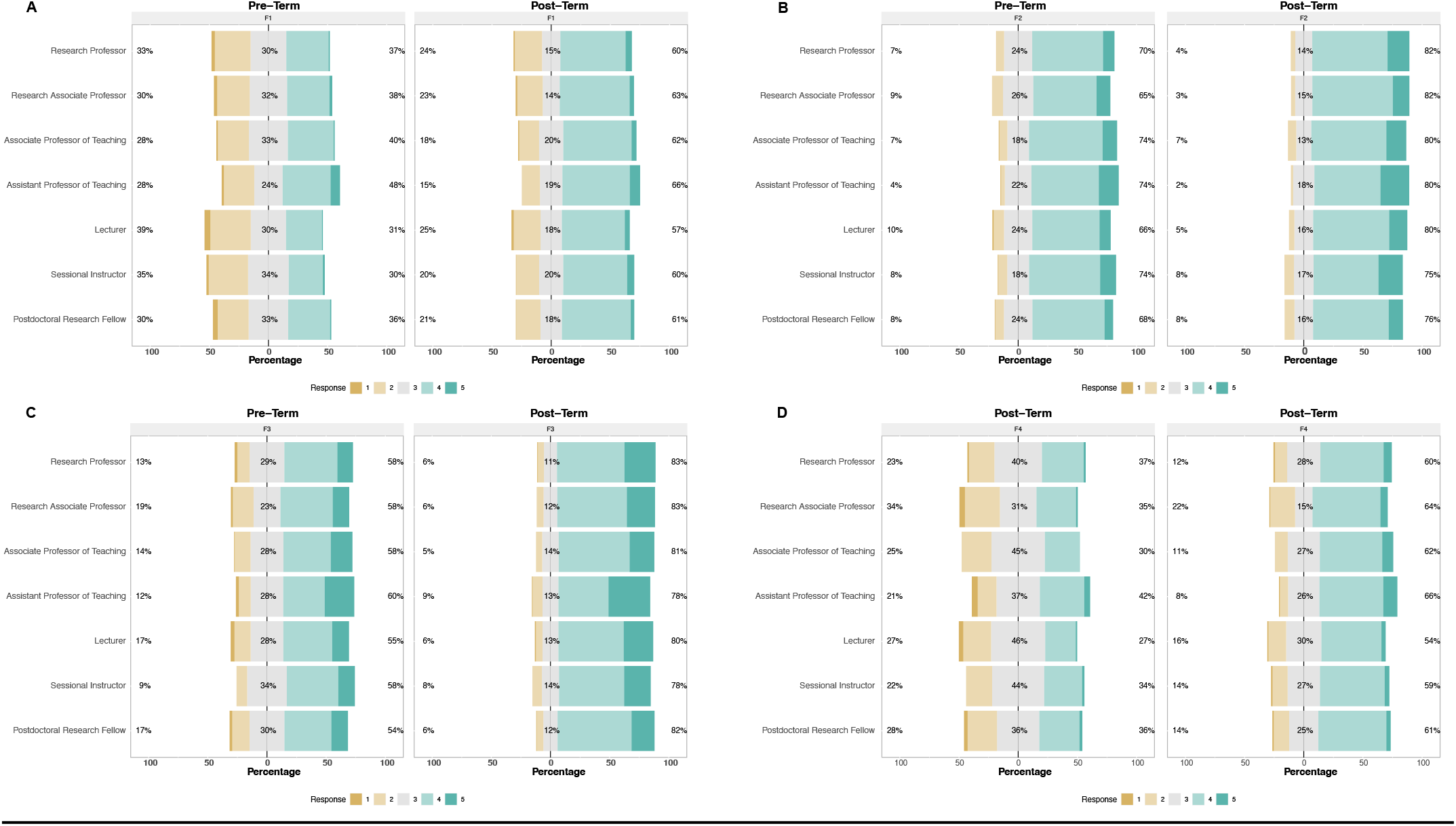
Pre- and post-term SAAB survey results of A) F1: *Decoding Primary Literature*, B) F2: *Interpreting Data*, C) F3: *Active Reading*, D) F4: *Visualization*. Response variables ranged from 1 “strongly disagree” at all to 5 “strongly disagree”, which represent the students self-identified confidence in their abilities in their skills before (pre-term) and after (post-term) taking BIOL140.

In addition, cohorts of Assistant Professors of Teaching and Sessional Instructors did not shift to the same degree as students of other instructor types in their ability to *actively read* primary literature (Fig. 2C, Suppl. Table 2). Interestingly, when we compared students’ pre-term responses to establish a baseline for observed gains across instructor types, we found higher confidence in cohorts of Assistant Professors of Teaching and Sessional Instructors than the other instructor types (Fig. 2C. Table 4). 58% of the students of Sessional Instructors and 60% of the students taught by the Assistant Professor of Teaching reported high or very high confidence in their ability to *actively read* scientific literature leading to a lower, non-significant gain to the end of the course (Fig. 2C).

**Table 4.**
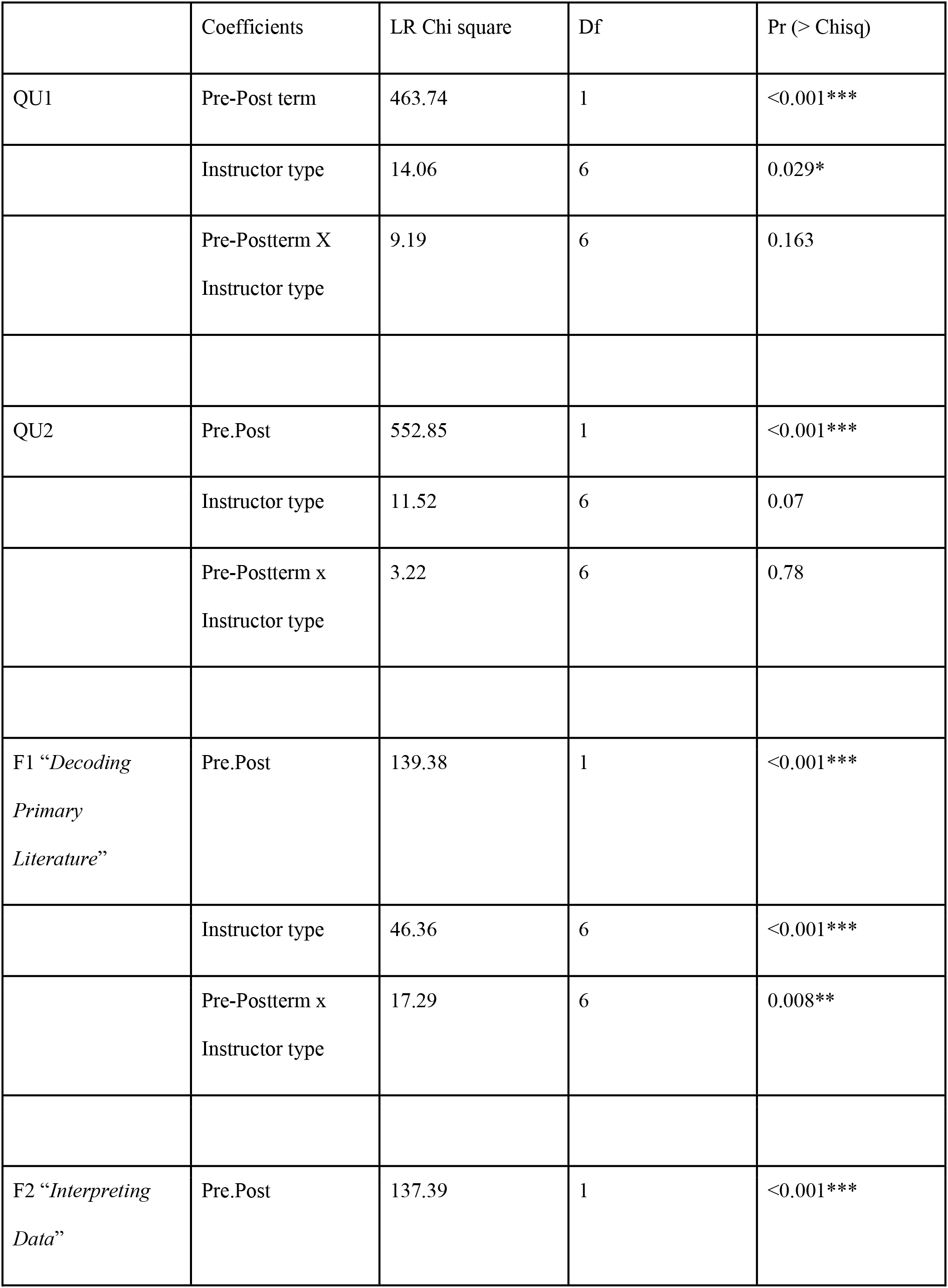
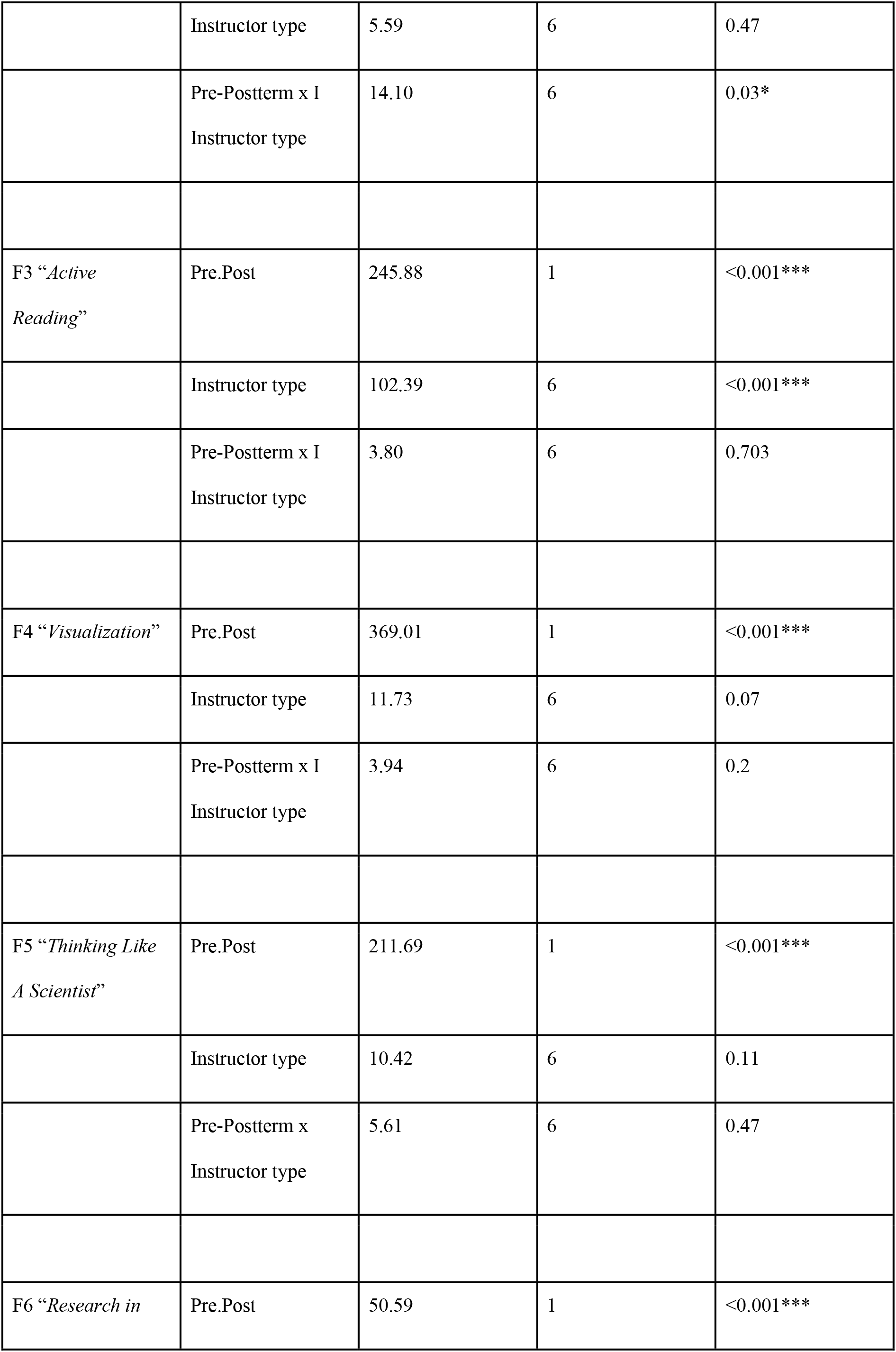
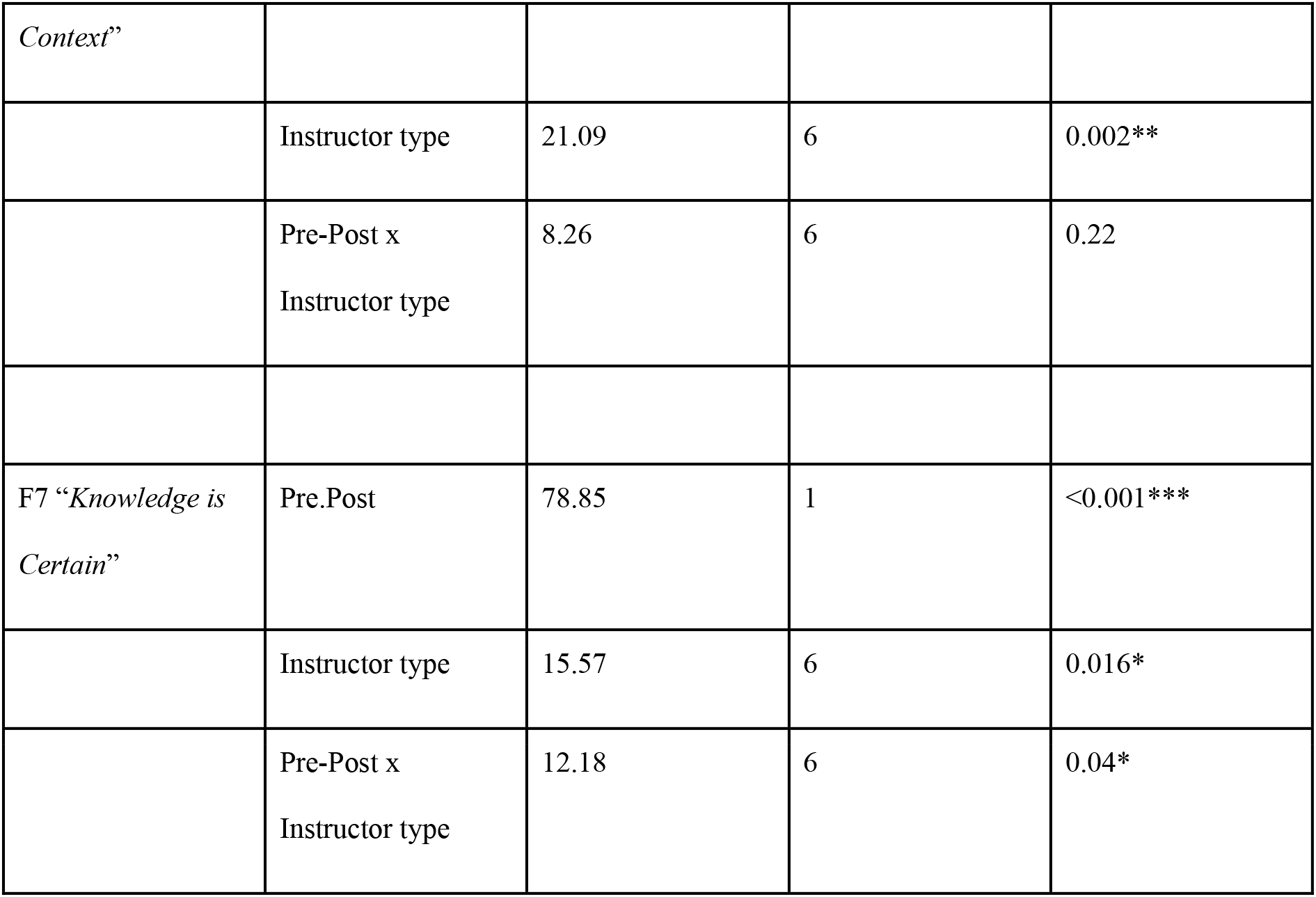
Ordinal logistic regression model results of survey using an analysis of variance (ANOVA). Df = degrees of freedom, pr = significance level of Chi Squared test. Asterix identifies significance levels with *** = p<0.001, ** = <0.01, * = <0.05.

While overall, students’ epistemological beliefs, or knowledge of the ‘*scientific publication process and the interpretation of data and forming conclusions*’ significantly increased from pre-term to post-term (p<0.0001, Table 2, Fig. 3A, B, C), several individual instructor types were unable to significantly shift their students’ epistemological beliefs (Table 4, Suppl. Table 1). Specifically, students of Postdoctoral Research Fellows, Associate Professors of Teaching, Assistant Professors of Teaching, and Research Associate Professors did not significantly improve from pre-to post-term surveys (p > 0.05, Table 4, Suppl. Table 2). In contrast, students of Research Professors, Lecturers, and Sessional Instructors significantly shifted from reporting “I am not sure” (3) to “I agree” (4) or “I strongly agree” (5) by 7% – 13%. (Fig. 3C. Table 4, Suppl. Table 1). Consistent with pre-term survey response data on skills and abilities, students of the Assistant Professor of Teaching came into the course with relatively sophisticated epistemological beliefs about science (Fig. 3C). Though shifts varied across instructor types, no significant differences were detected between students taught by instructors of different ranks at the end of term (Table 4).

**Figure 3.**
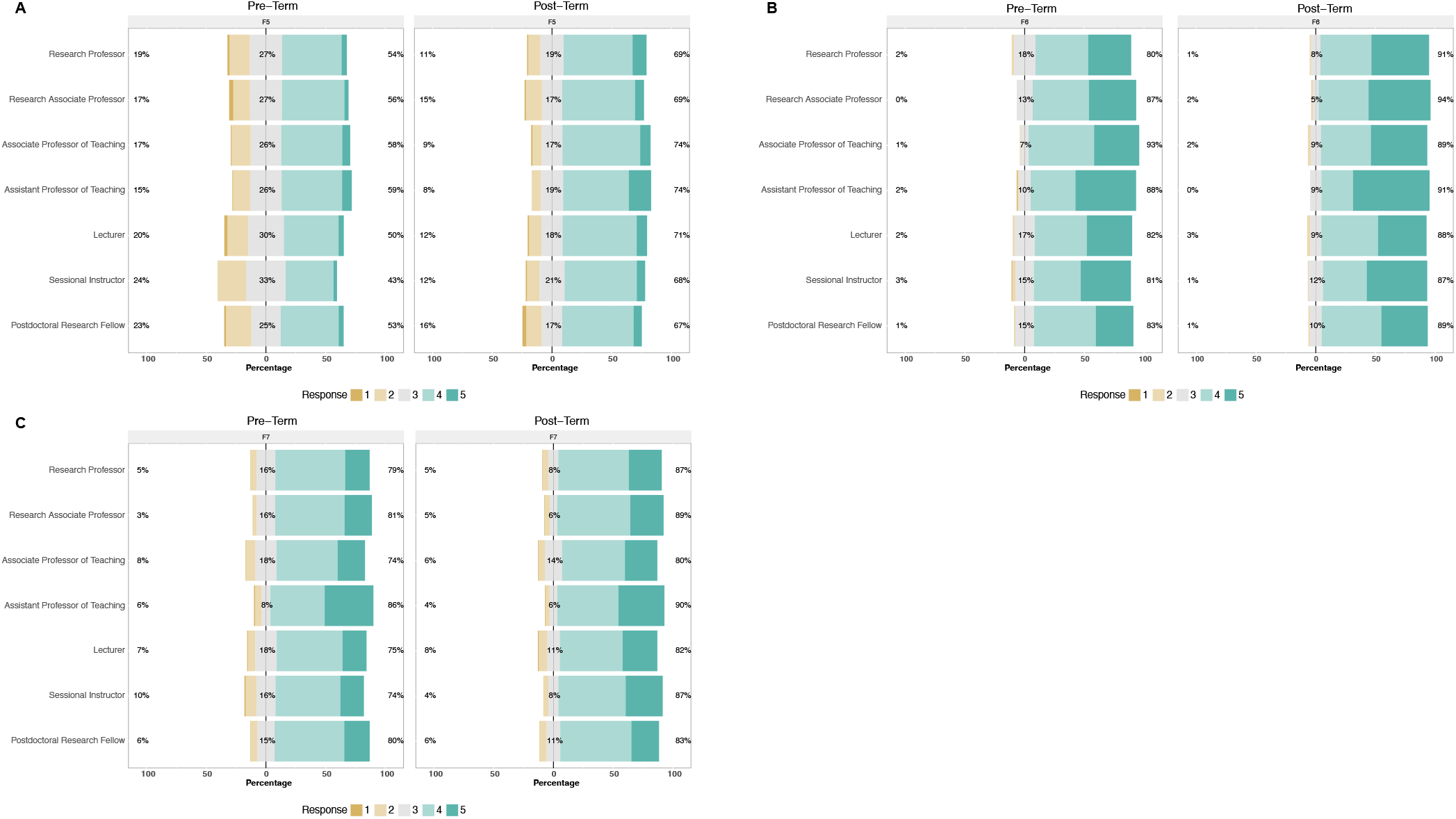
Pre- and post-term surveys SAAB result of A) F5: *Thinking Like A Scientist*, B) F6: *Research in Context*, C) F7: *Knowledge is Certain.* Response variables ranged from 1 being not confident at all to 5 being very confident, which represent the students self-identified confidence in their abilities in their skills before (pre-term) and after (post-term) taking BIOL140.

## Discussion

In the present study, we extended the utility of the CREATE pedagogy to a large, multi-section, team-taught first-year undergraduate biology course. We showed that it was successfully implemented in the online learning environment. We demonstrated that CREATE outcomes did not depend on instructor rank, training, and/or teaching experience but rather were achieved through the collaborative work of instructors who met regularly and worked as a learning community. We observed positive shifts in students’ confidence in their science skills and abilities across multiple sections taught by different instructors of a single course. We showed that instructors using CREATE online achieved comparable outcomes. Overall, our results expand on the current and growing body of literature demonstrating that the CREATE pedagogy fosters student efficacy by engaging students in the practices of scientists and extends these outcomes to the virtual classroom.

CREATE has previously been shown to increase students’ self-assessed ability to successfully carry out a range of skills required to comprehend primary scientific literature (8, 11, 14). By engaging students in an iterative practice of synthesizing complex information, thinking critically through study design, and interpreting the findings of published science, CREATE helps students cultivate the skills of scientists (11, 15, 16). Our findings underscore these studies and extend the use of the student-centered activities defining CREATE pedagogy (i.e. concept mapping, article and figure annotation, and data transformation) to the online teaching and learning environment.

We show that instructors of various ranks and levels of teaching experience can achieve similar student outcomes in the affective domain when implementing the evidence-based teaching practice CREATE with no formal training. We observed no differences in student outcomes between tenured and untenured instructors. Kenyon et al. (8) hypothesized that professional status may influence student outcomes as tenured faculty may have “more flexibility in course selection or design” than untenured faculty. While this may be true if different parameters constrained individual instructors, our data indicate that the pedagogy is versatile when implemented by instructors of various ranks and when working with a comparable degree of flexibility. Although individual categories of questions in this study revealed differences between instructors, such differences can be explained by pre-term survey results, which show differences across sections in students coming into the course. For example, we found significant differences between instructors in the question categories ‘*F1: Decoding Primary Literature*’ and ‘*F3: Active Reading,*’; however, these were due to differences in student’s self-assessed abilities pre-term. These findings demonstrate that the CREATE approach can shift students’ self-assessed beliefs in their ability to think and do science. These results argue that neither professional status nor teaching experience affects student outcomes but do not pinpoint how instructors achieved comparable outcomes. Previous research has elucidated the value of alignment between instructor intentions, defined as “instructors’ goals for carrying out a specific task with students” and instructor support, described as “what instructors say to students during interactions” (17). In the present study, the teaching team explicitly discussed their shared intentions each week. Still, individual instructor supports likely varied across sections, given the range of teaching and research experiences that different instructors brought to the course. Future work may characterize instruction in different learning environments using instruments like the Classroom Discourse Observation Protocol (18) and explore how instructor talk influences student outcomes across instructor types (19).

Unlike previous studies in which newly trained faculty achieved positive student gains and outcomes after attending a CREATE workshop facilitated by experts, none of the instructors in this study received formal training before the start of their CREATE course (8, 12, 13). Due to the immediate shift to remote teaching and learning during the COVID-19 pandemic, this course was quickly redesigned for remote teaching and learning. Instructors needed to have the opportunity to receive formal training in new pedagogy or online tools. However, we show that faculty new to teaching CREATE and who are also new to online teaching and learning can successfully increase students’ efficacy and epistemological beliefs about science.

### An instructor learning community supports the adoption of evidence-based practice

We attribute the successful implementation of CREATE here to our instructors’ commitment to a learning community. We define an instructor learning community as a cross-disciplinary group of various instructor types which collaborates as an instructional team throughout the term but differentiates between our *instructor* learning community and more commonly referenced *faculty* learning community as not all participants were faculty members (20). Instructors actively participated in CREATE learning activities during our weekly community meetings. The meetings provided a space for instructors to discuss challenges, highlight successes, and build confidence before taking the risks involved in implementing new pedagogy. Previous studies have found that different instructors make different decisions on their instructional strategies based on their attitudes rather than pedagogical evidence (21, 22). In the present study, the instructor learning community was facilitated by a faculty member that centered discussions on pedagogical evidence and shared goals but also created space for individuals to reflect on and share their attitudes and beliefs about teaching and learning. Facilitation was guided by the principles of self-determination theory in which instructors cultivated *competence* by piloting CREATE activities, *relatedness* by connecting over shared goals and challenges, and *autonomy* by bringing themselves and their own choices into weekly meetings (23). Our results indicate that instructor learning communities may attenuate barriers to pedagogical change within departments by providing inclusive opportunities to engage instructors of various ranks in shared goals and discussion on teaching and learning. Unlike many professional development opportunities that facilitate the renewal of teaching practices, such as workshops and seminars, which require voluntary and additional time commitments that impede the introduction of new pedagogy, our instructor learning community was an integral part of the course.

### The affective domain of learning as a course design feature for high-impact teaching

The affective domain of learning is concerned with the feelings that arise throughout the learning process, which influence learning outcomes. It is distinguished from the cognitive domain of learning in that feelings and attitudes define it as opposed to knowledge and intellectual skills (24). This study analyzed student self-assessment data to explore the affective domain of learning, precisely the construct of students’ science efficacy. Increasing students’ confidence in their ability to *do science* or their science efficacy is associated with numerous positive outcomes (25, 26). Self-efficacious students are intrinsically motivated, more likely to challenge themselves, able to recover from failure and setbacks, and more likely to achieve their goals. Increases in student efficacy have even been shown to close gaps and improve the academic performance of students historically underserved in science (27).

High-impact teaching activities that promote collaboration and cultivate community have been shown to facilitate short and long-term outcomes such as increased efficacy and persistence in STEM (25, 28–30). Students in this study worked in fixed small groups on CREATE activities throughout the term and informal feedback from students corroborates the idea that collaboration in the form of group work was an integral part of the course that likely contributed to a sense of belonging and connectedness, which may have in turn enhanced science efficacy. While this study extends the utility of the CREATE pedagogy to both the online environment and a diversity of instructors, our work needs to elucidate the exact mechanism through which CREATE increases students’ science efficacy. Future work characterizing the students’ experience of CREATE activities would increase our understanding of how the pedagogy supports the affective domain of learning.

As we move beyond many of the challenges that the COVID-19 pandemic initially posed to post-secondary education, we carry with us valuable lessons learned about the noncognitive factors that contribute to student success and hopefully a heightened sense of empathy for our students. This is particularly important in the STEM fields where the affective domain of learning has traditionally garnered less attention than the cognitive domain yet when given sufficient attention, has been shown to help all STEM students thrive, especially those with visible or invisible disabilities and those from historically underrepresented backgrounds that may be hesitant about their social belonging (29, 31–34).

## ACKNOWLEDGMENTS

We want to acknowledge that this study was conducted on the traditional, ancestral, and unceded territory of the Musqueam (xʷməθkʷə^y̓^ əm) people, which has been a place of learning for many generations before the University of British Columbia was built on these lands. We thank the University of British Columbia and the Biology Program for their support. We thank all instructors who taught this course: Chin Sun, Andrew Trites, Brett Couch, Brittany Carr, Curtis Suttle, George Haugh, Jennifer Klenz, Ljerka Kunst, Matt Ramer, Phil Matthews, Sarah Moore, Stella Lee, Yuelin Zhang, Catlin Donelly. Darren Irwin, Geoff Wasteneys, Jackie Dee, Naomi Fast, Skye Butterson, Trish Schulte, and Joerg Bohlmann. We also extend our thanks to Warren Code for their valuable input and insightful suggestions, which greatly enhanced the quality of this study. This study was supported by the UBC Vancouver Advancing Education Renewal Fund. The collection of student responses in BIOL140 was approved by the University of British Columbia’s Behavioural Research Ethics Board (H21-03606)

## Appendix

**Supplementary Table 1.** Cumulative link model outputs for testing for significant effects between terms. SE = standard error, df = degrees of freedom, p valie = significance level

| Question Group | Parameter contrasts | Estimates | SE | df | P value |
| --- | --- | --- | --- | --- | --- |
| QU1 | Pre Term 1 –<br>Pre Term 2 | -0.437 | 0.220 | Inf | 0.193 |
|  | Post Term 1 –<br>Post Term 2 | -0.238 | 0.217 | Inf | 0.691 |
| QU2 | Pre Term 1 –<br>Pre Term 2 | -0.464 | 0.197 | Inf | 0.085 |
|  | Post Term 1 –<br>Post Term 2 | -0.212 | 0.194 | Inf | 0.695 |
| F1 “*Decoding Primary Literature*” | Pre Term 1 –<br>Pre Term 2 | 0.117 | 0.109 | Inf | 0.709 |
|  | Post Term 1 –<br>Post Term 2 | 0.104 | 0.109 | Inf | 0.779 |
| F2 “*Interpreting Data*” | Pre Term 1 –<br>Pre Term 2 | 0.324 | 0.137 | Inf | 0.084 |
|  | Post Term 1 –<br>Post Term 2 | 0.268 | 0.137 | Inf | 0.207 |
| F3 “*Active Reading*” | Pre Term 1 –<br>Pre Term 2 | 0.783 | 0.088 | Inf | 0.812 |
|  | Post Term 1 –<br>Post Term 2 | 0.022 | 0.087 | Inf | 0.994 |
| F4 “*Visualization*” | Pre Term 1 –<br>Pre Term 2 | 0.443 | 0.159 | Inf | **0.029** |
|  | Post Term 1 –<br>Post Term 2 | 0.364 | 0.159 | Inf | 0.107 |
| F5 “*Thinking Like A Scientist*” | Pre Term 1 –<br>Pre Term 2 | -0.146 | 0.120 | Inf | 0.615 |
|  | Post Term 1 –<br>Post Term 2 | 0.028 | 0.120 | Inf | 0.995 |
| F6 “*Research in Context*” | Pre Term 1 –<br>Pre Term 2 | -0.107 | 0.135 | Inf | 0.858 |
|  | Post Term 1 –<br>Post Term 2 | 0.129 | 0.137 | Inf | 0.784 |
| F7 “*Knowledge is Certain*” | Pre Term 1 –<br>Pre Term 2 | -0.195 | 0.107 | Inf | 0.264 |
|  | Post Term 1 –<br>Post Term 2 | 0.019 | 0.107 | Inf | 0.997 |

**Supplementary Table 2.** Model comparison using the likelihood ratio test of cumulative link models. Link = threshold; logit distribution = equidistant. Asterix identifies significance levels with *** = p<0.001, ** = <0.01, * = <0.05.

|  |  |  |  |  |  |  |
| --- | --- | --- | --- | --- | --- | --- |
| QU1 | model.fixed : Likert score ~ Pre.Post + Instructor.type + Pre.Post:Instructor.type<br>Model: Likert score ~ Pre.Post + Instructor.type + Pre.Post:Instructor.type + (1\|Student.Code) |  |  |  |  |  |
|  | No. par | AIC | logLik | LR.stat | df | Pr(>Chisp) |
| model.fixed | 15 | 2792.7 | -1381.4 |  |  |  |
| model | 16 | 27.13.5 | -1340.7 | 81.25 | 1 | <0.001*** |
| QU2 | model.fixed : Likert score ~ Pre.Post + Instructor.type + Pre.Post:Instructor.type<br>Model: Likert score ~ Pre.Post + Instructor.type + Pre.Post:Instructor.type + (1\|Student.Code) |  |  |  |  |  |
|  | No. par | AIC | logLik | LR.stat | df | Pr(>Chisp) |
| model.fixed | 15 | 2552.9 | -1261.5 |  |  |  |
| model | 16 | 2525.1 | -1246.6 | 29.78 | 1 | <0.001*** |
| F1<br>“Decoding<br>Primary<br>Literature” | model.fixed : Likert score ~ Pre.Post + Instructor.type + Pre.Post:Instructor.type<br>Model: Likert score ~ Pre.Post + Instructor.type + Pre.Post:Instructor.type + (1\|Student.Code) |  |  |  |  |  |
| model.fixed | 15 | 7743.2 | -3856.6 |  |  |  |
| model | 16 | 7381.9 | -3675.0 | 363.32 | 1 | <0.001*** |
| F2<br>“*Interpreting Data*” | model.fixed : Likert score ~ Pre.Post + Instructor.type + Pre.Post:Instructor.type<br>Model: Likert score ~ Pre.Post + Instructor.type + Pre.Post:Instructor.type + (1\|Student.Code) |  |  |  |  |  |
| model.fixed | 15 | 12857 | -6413.4 |  |  |  |
| model | 16 | 12132 | -6.049.8 | 727.37 | 1 | <0.001*** |
| F3 “*Active Reading*” | model.fixed : Likert score ~ Pre.Post + Instructor.type + Pre.Post:Instructor.type<br>Model: Likert score ~ Pre.Post + Instructor.type + Pre.Post:Instructor.type + (1\|Student.Code) |  |  |  |  |  |
| model.fixed | 15 | 17480 | -8724.8 |  |  |  |
| model | 16 | 17461 | -8714.7 | 20.39 | 1 | <0.001*** |
| F4<br>“*Visualization*” | model.fixed : Likert score ~ Pre.Post + Instructor.type + Pre.Post:Instructor.type<br>Model: Likert score ~ Pre.Post + Instructor.type + Pre.Post:Instructor.type + (1\|Student.Code) |  |  |  |  |  |
| model.fixed | 15 | 10924 | -5447.1 |  |  |  |
| model | 16 | 10145 | -5056.5 | 781.22 | 1 | <0.001*** |
| F5 “*Thinking Like a Scientist*” | model.fixed : Likert score ~ Pre.Post + Instructor + Pre.Post:Instructor<br>Model: Likert score ~ Pre.Post + Instructor + Pre.Post:Instructor + (1\|Student.Code) |  |  |  |  |  |
| model.fixed | 15 | 14682 | -7325.8 |  |  |  |
| model | 16 | 14232 | -7099.8 | 451.98 | 1 | <0.001*** |
| F6<br>*“Research in Context”* | model.fixed : Likert score ~ Pre.Post + Instructor.type + Pre.Post:Instructor.type<br>Model: Likert score ~ Pre.Post + Instructor.type + Pre.Post:Instructor.type + (1\|Student.Code) |  |  |  |  |  |
| model.fixed | 15 | 9189.6 | -4579.8 |  |  |  |
| model | 16 | 8901.7 | -44.34.8 | 289.94 | 1 | <0.001*** |
| F7<br>*“Knowledge is Certain”* |  |  |  |  |  |  |
| model.fixed | 15 | 20081 | -10025.7 |  |  |  |
| model | 16 | 19348 | -9658.2 | 735.06 | 1 | <0.001* |

**Supplementary Table 2.**
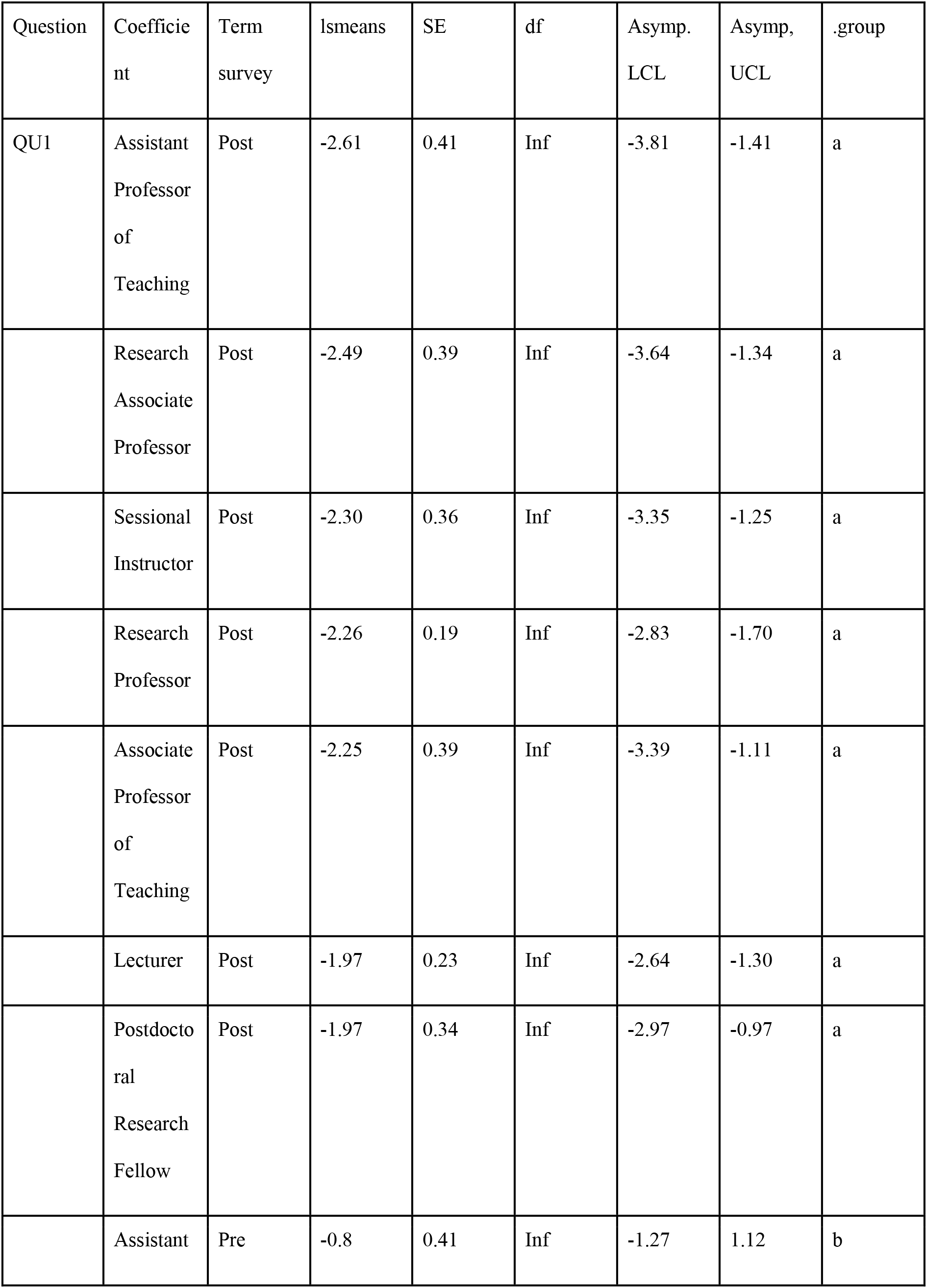
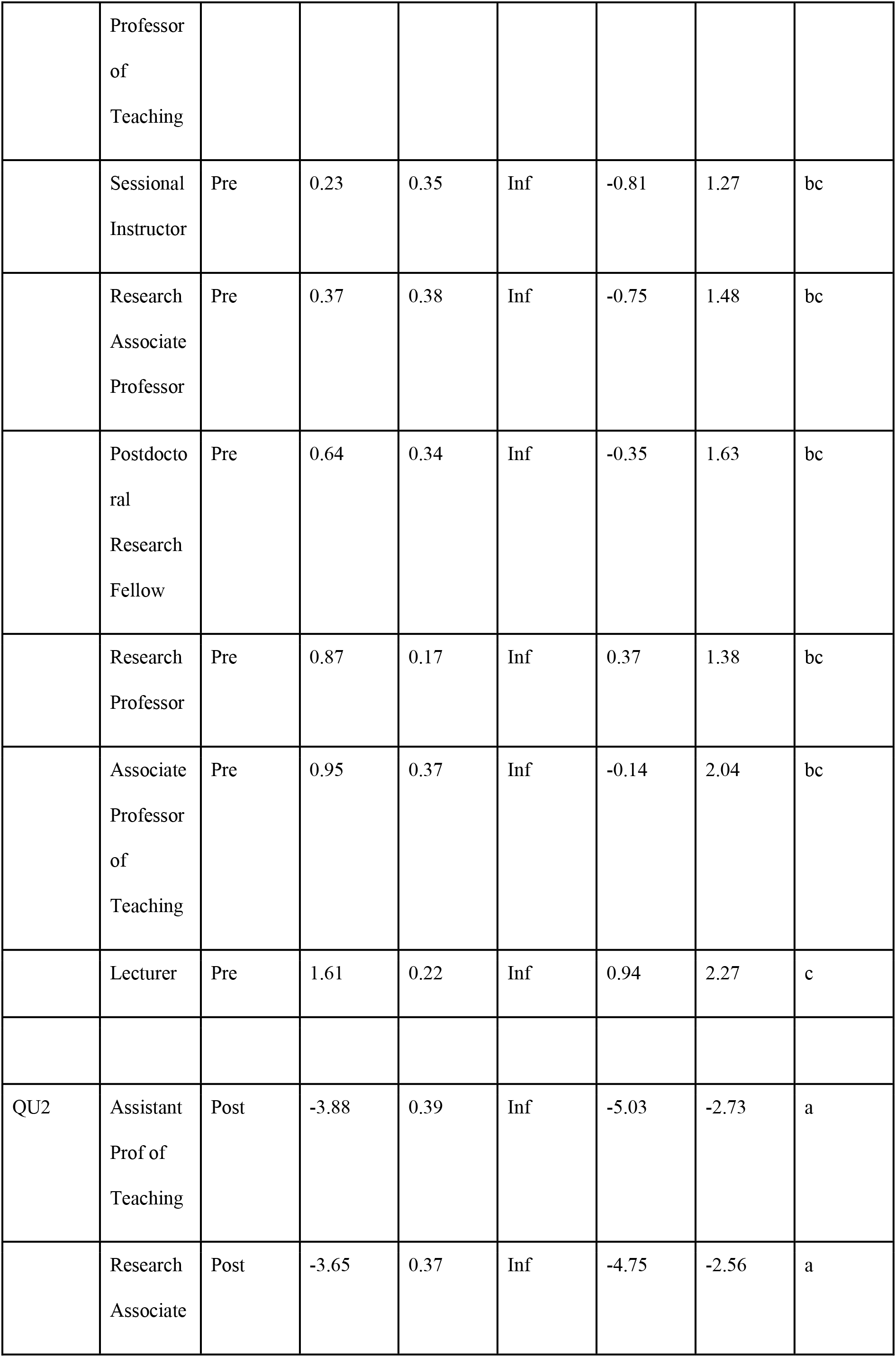
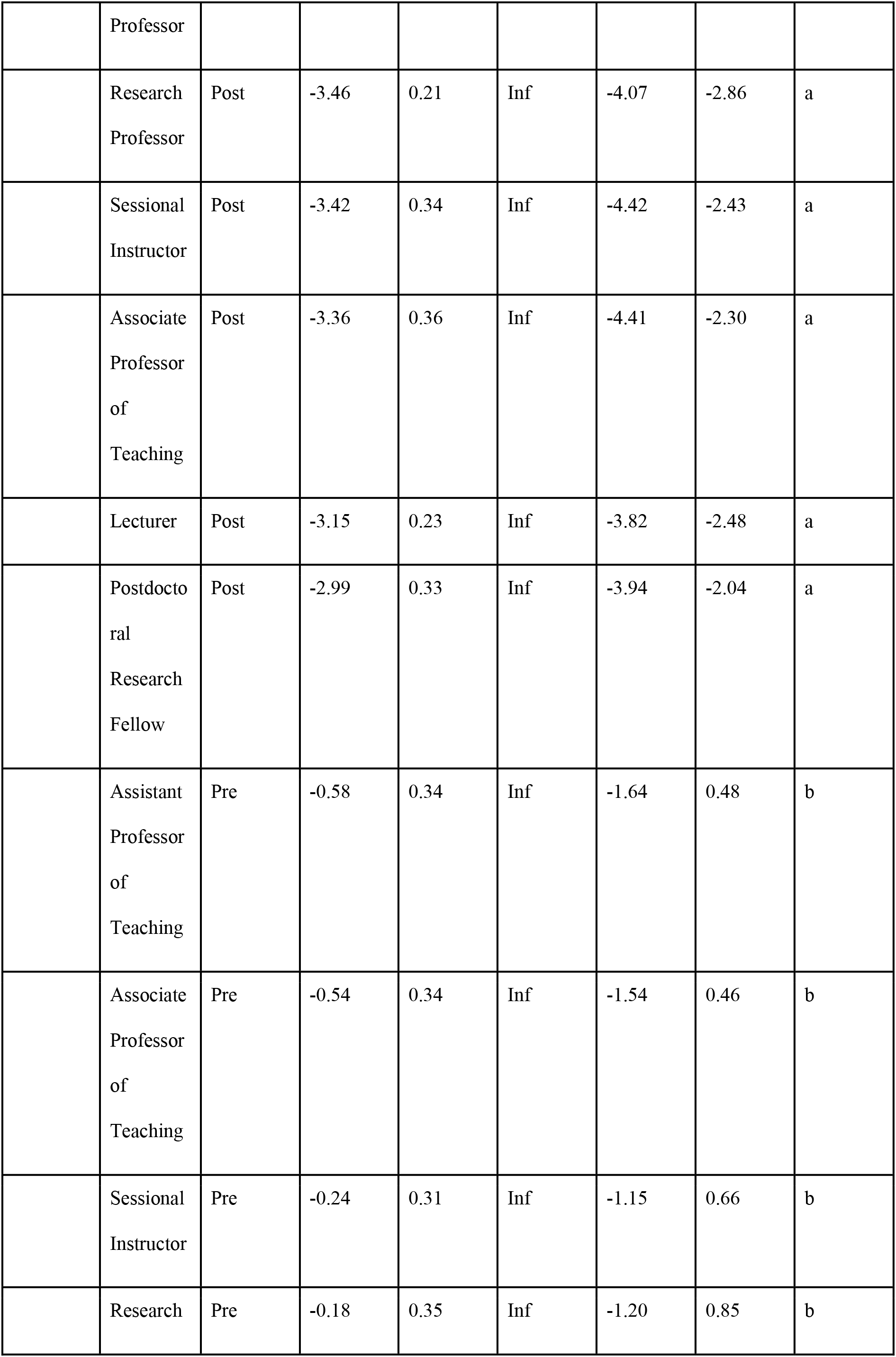
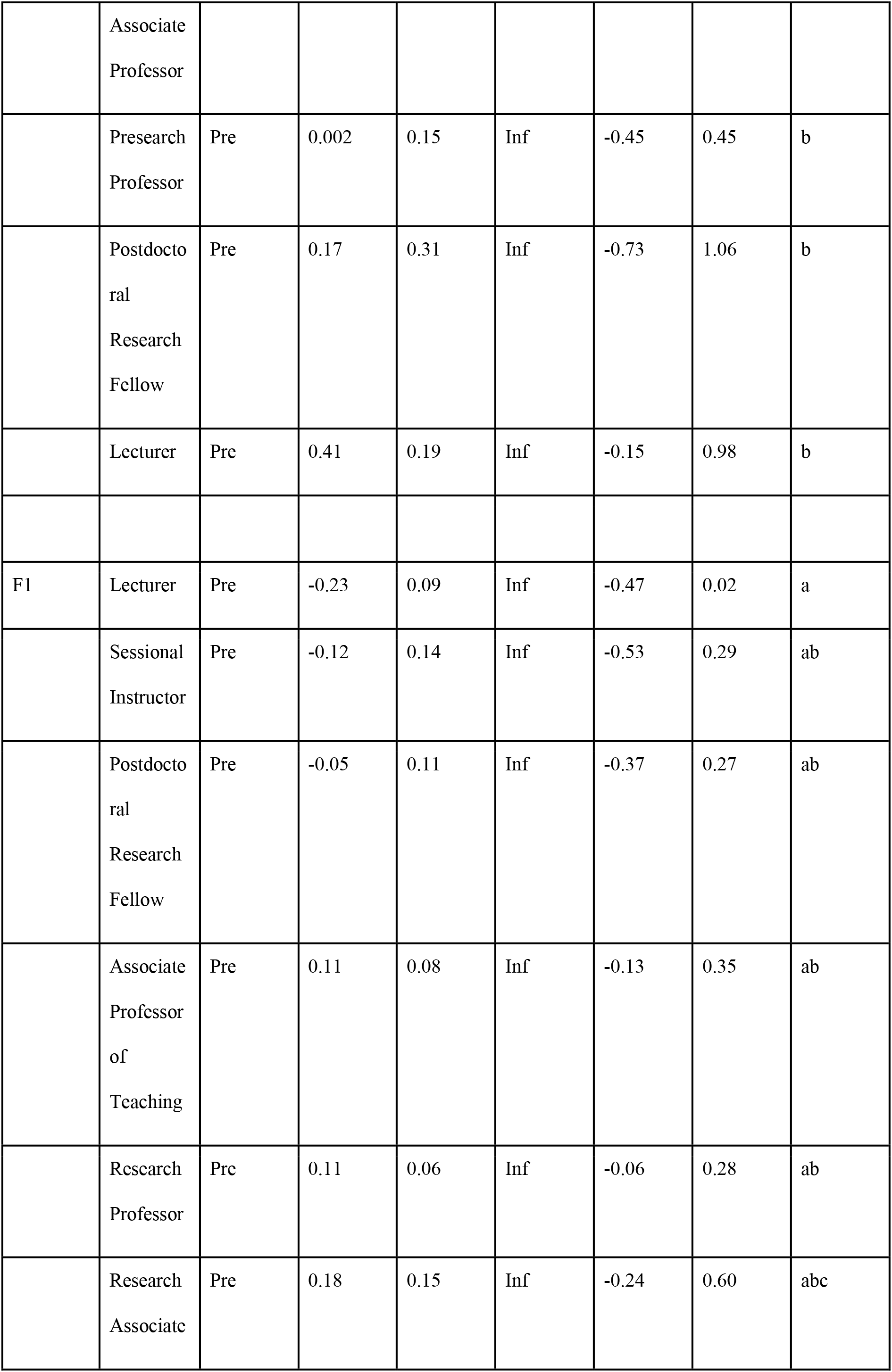
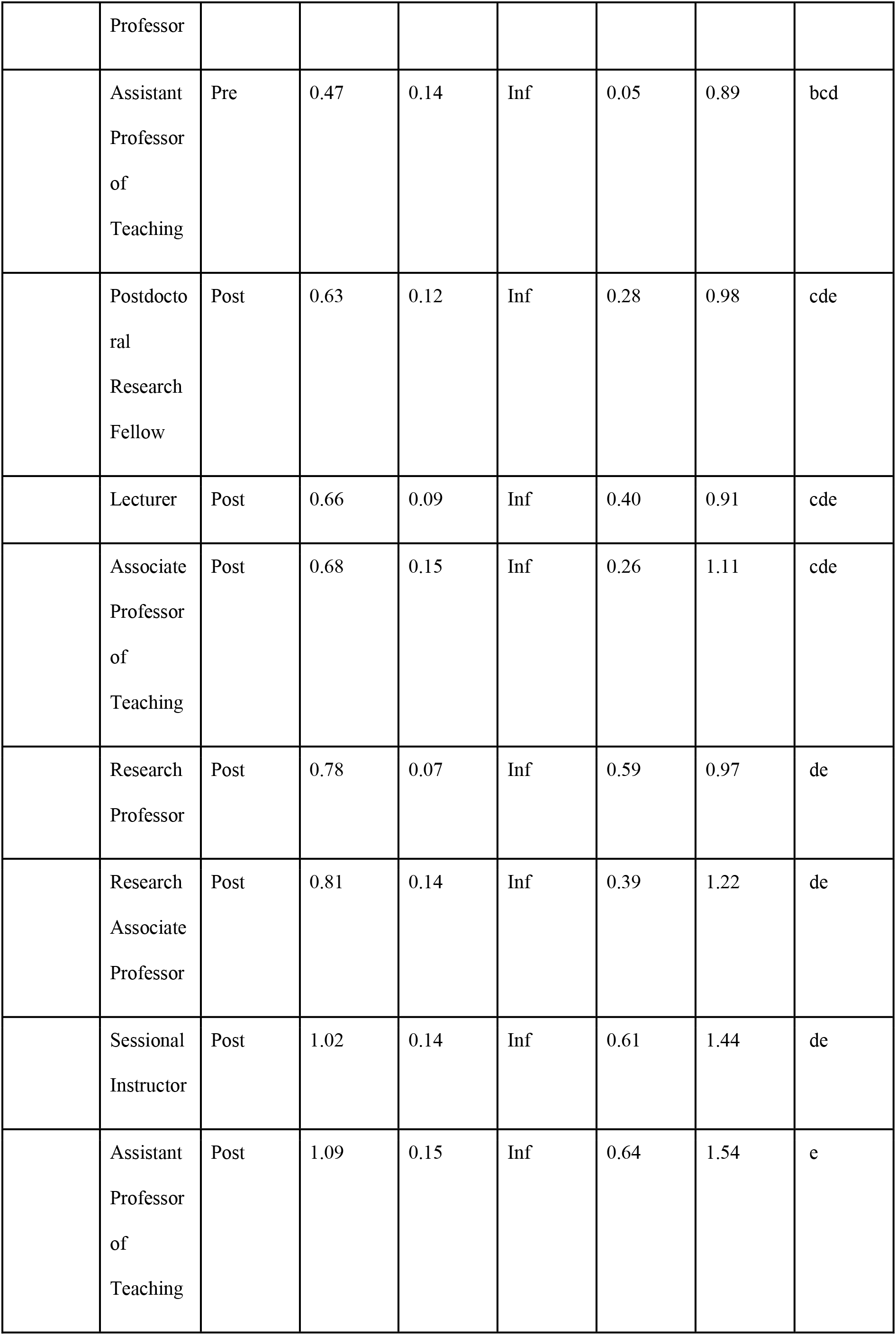
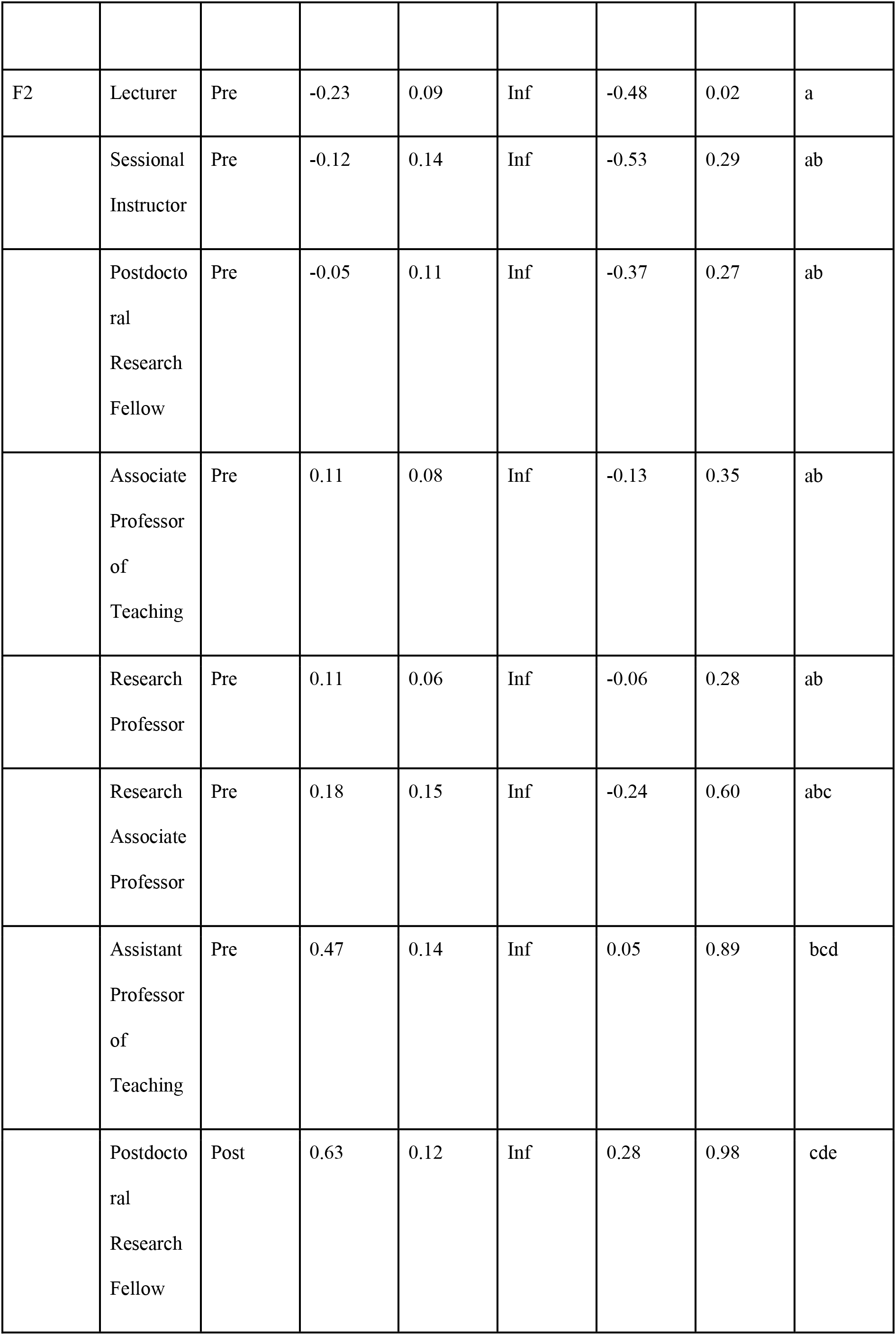
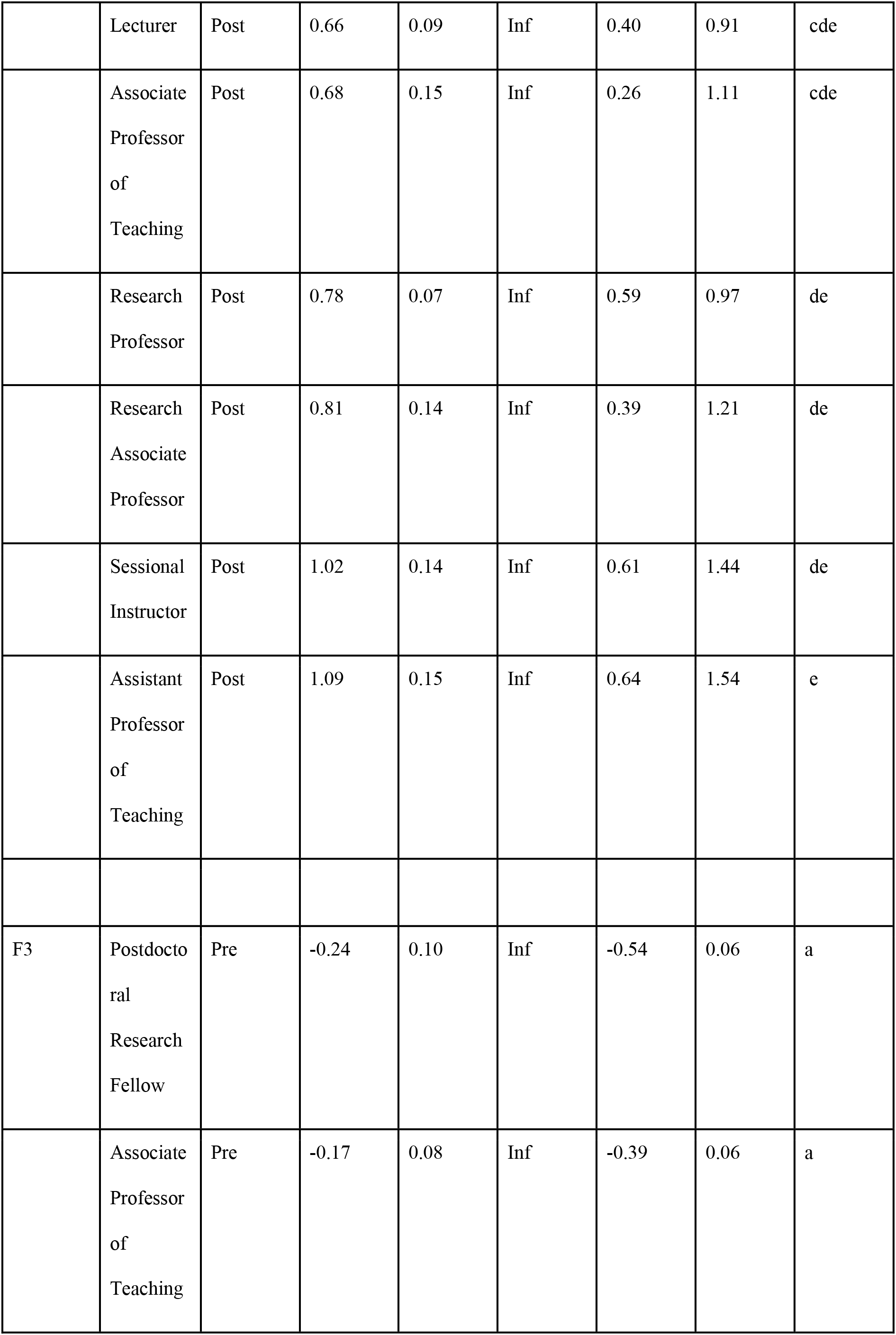
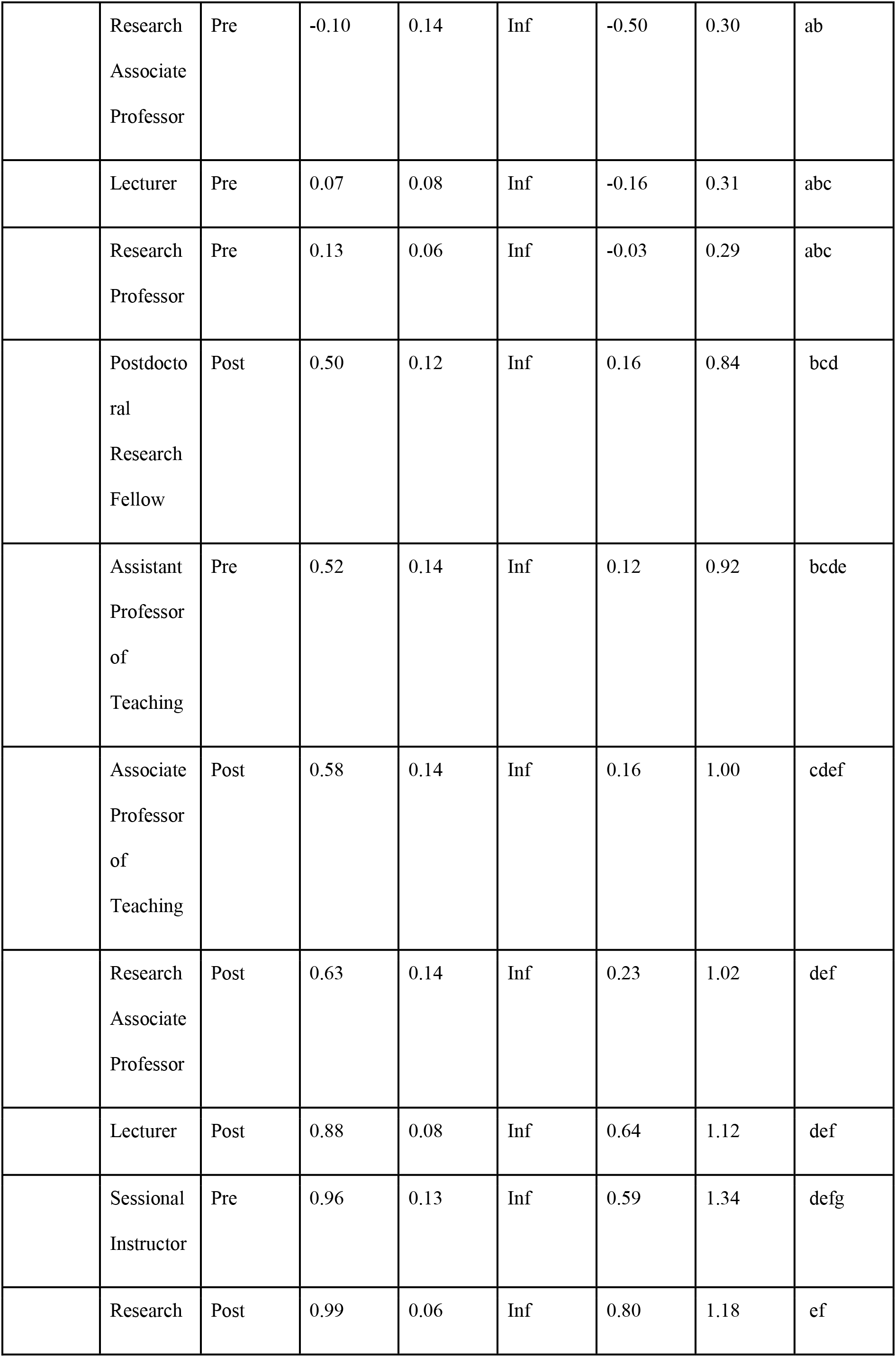
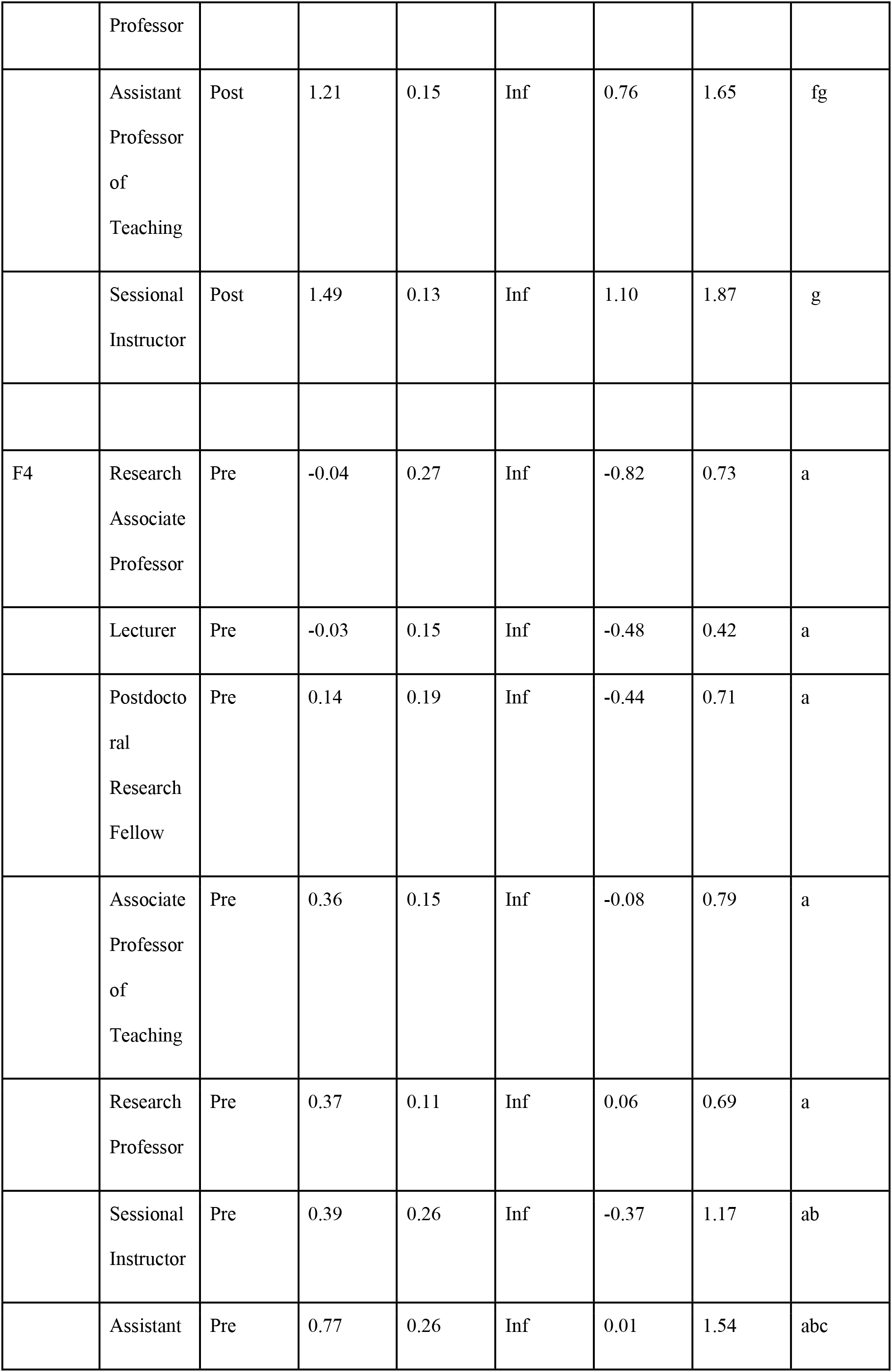
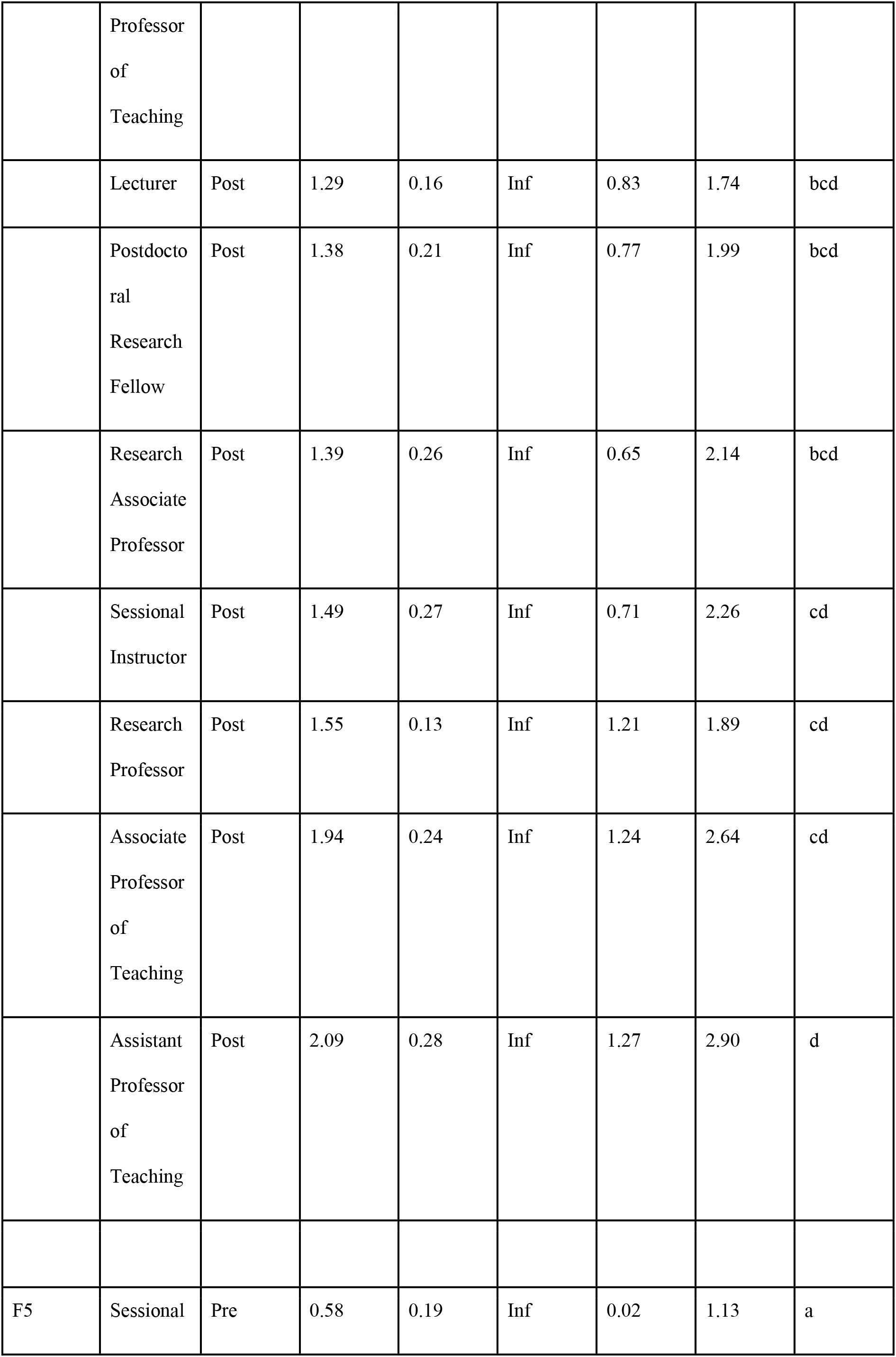
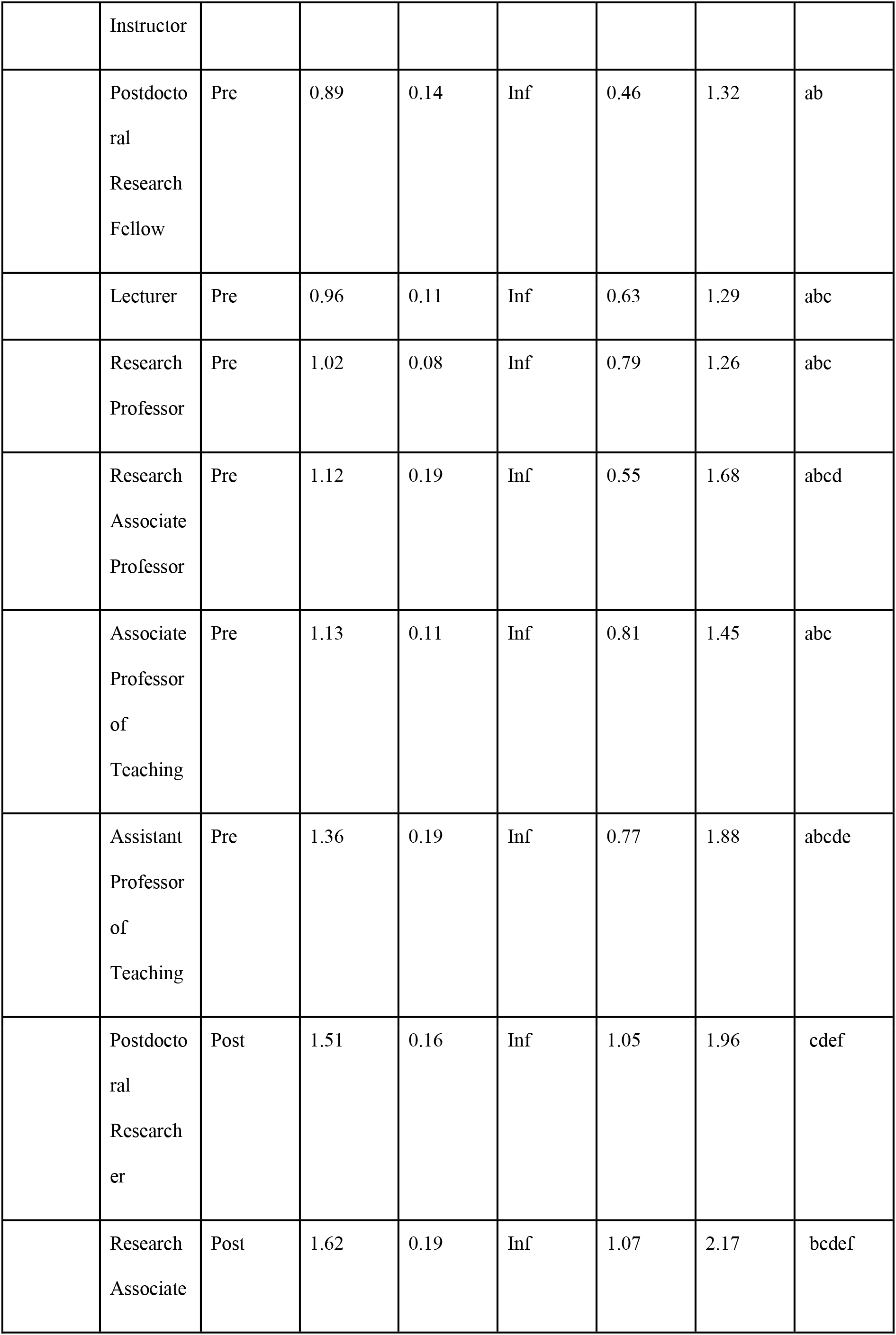
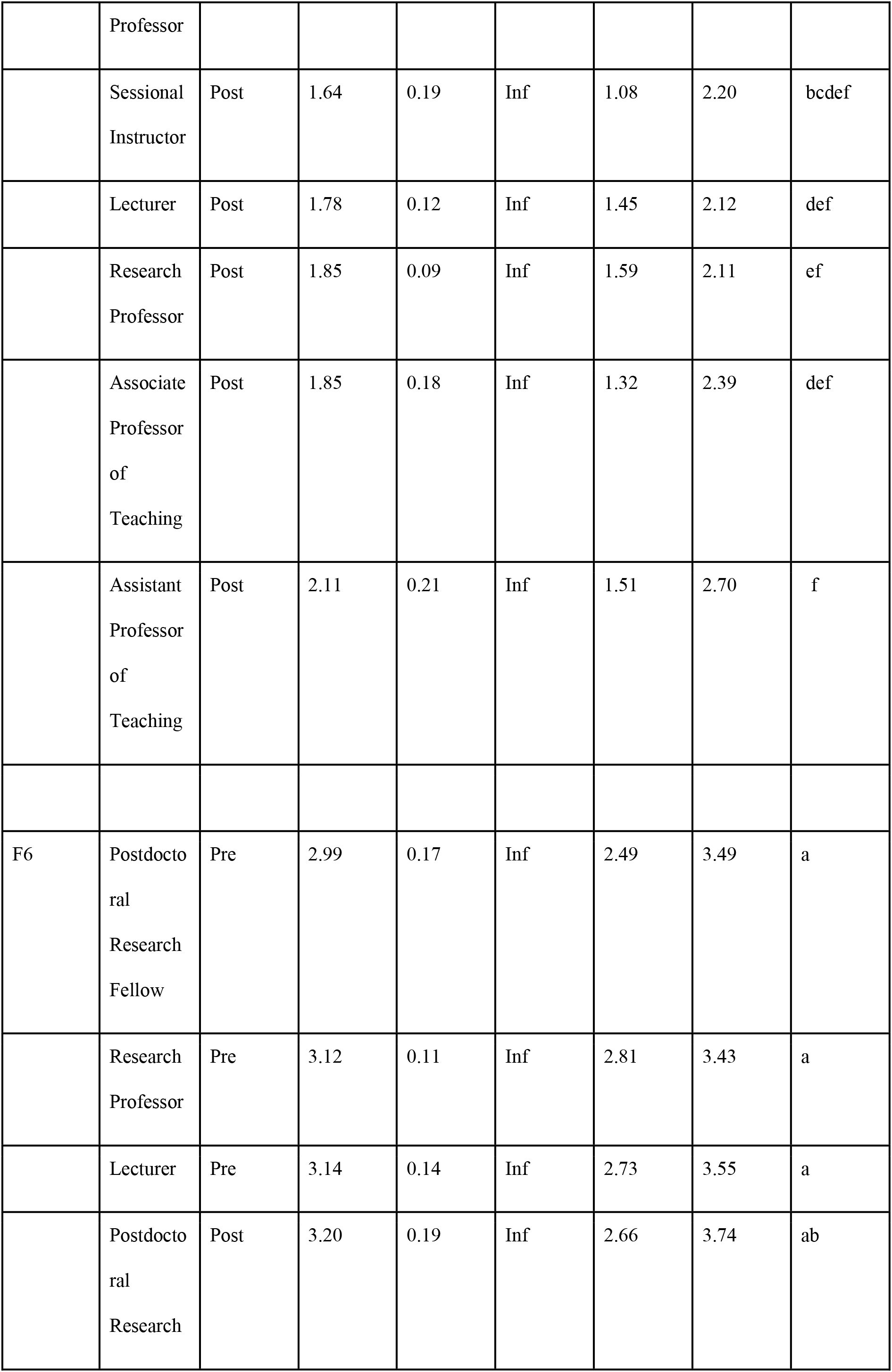
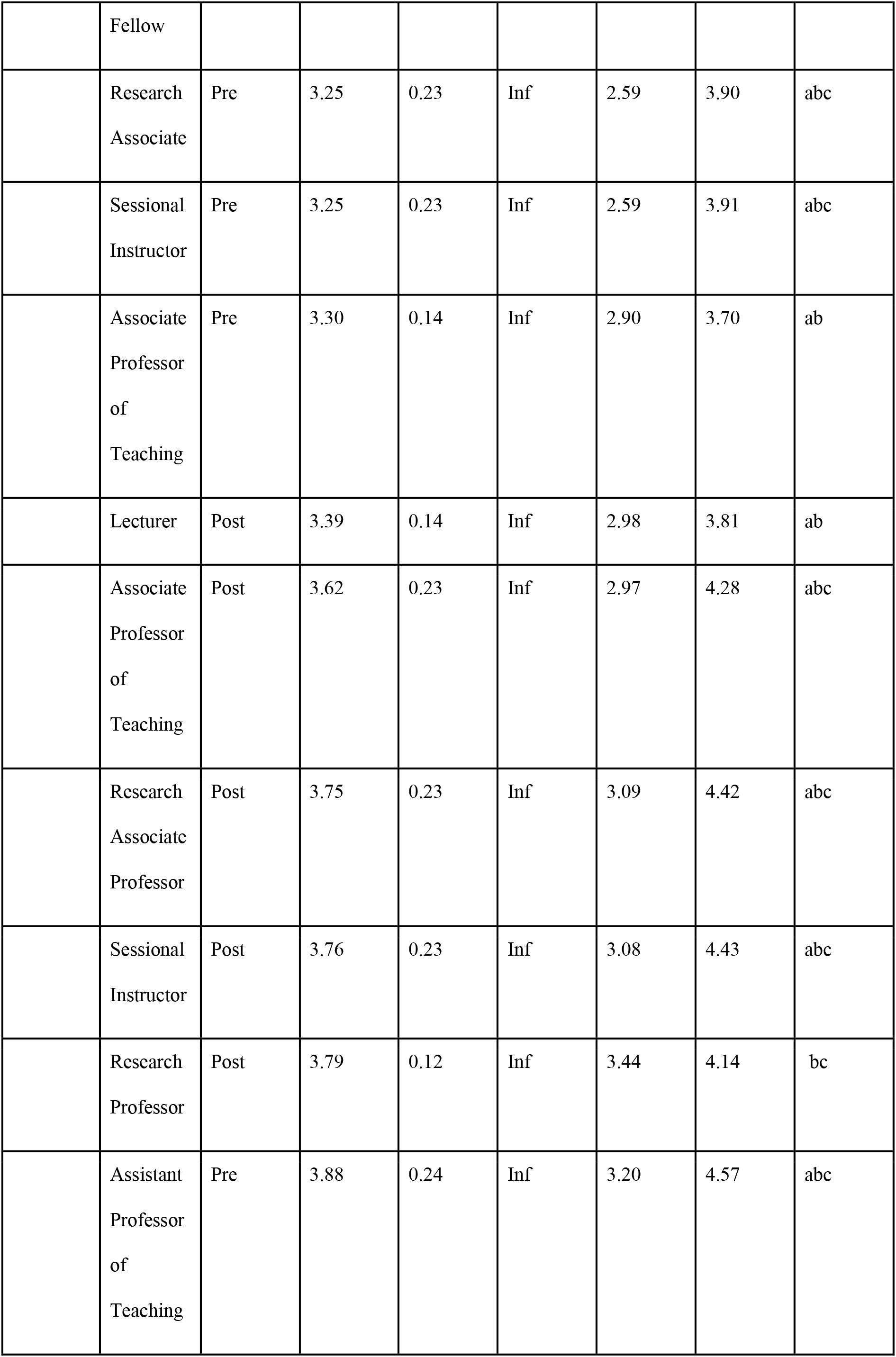
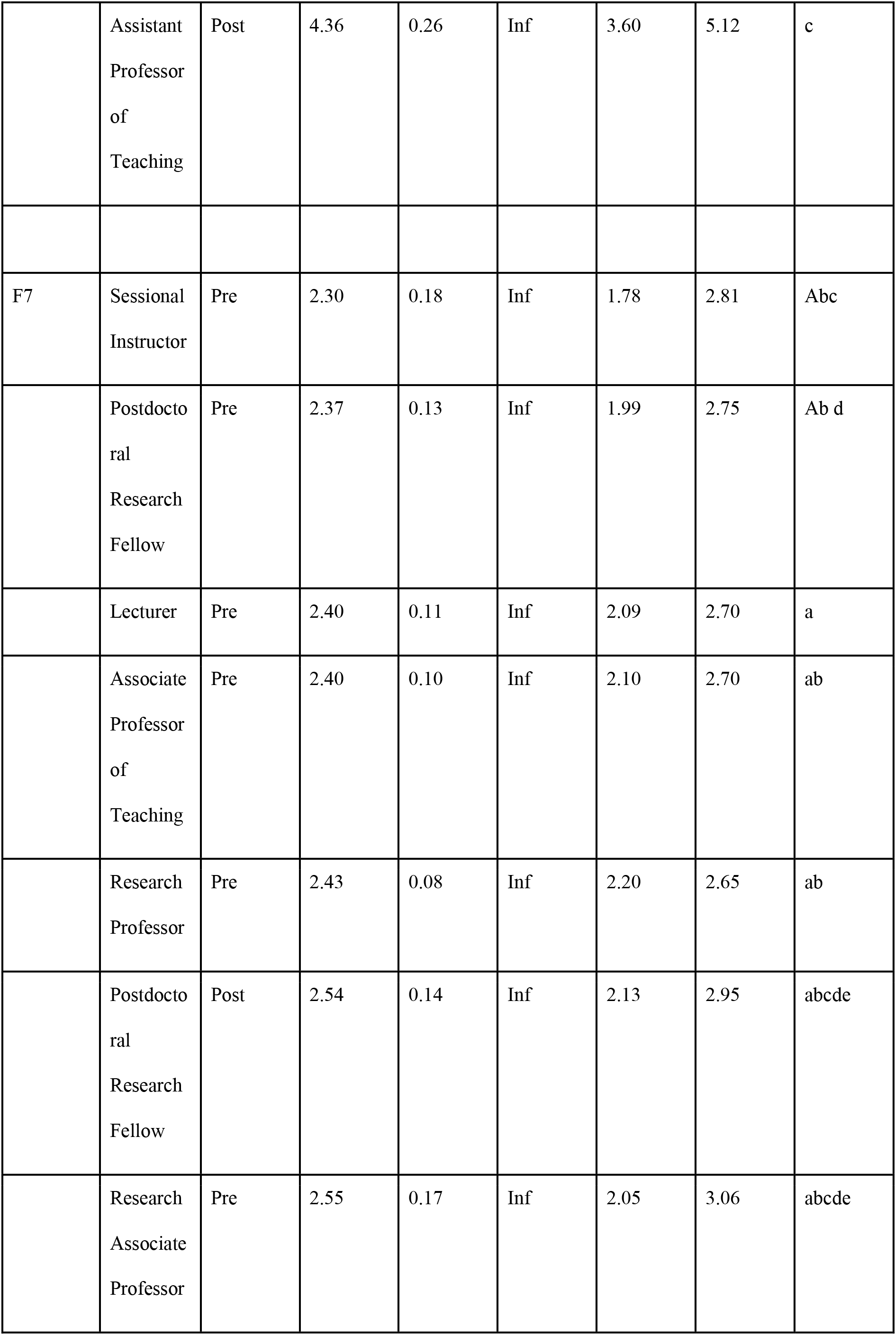
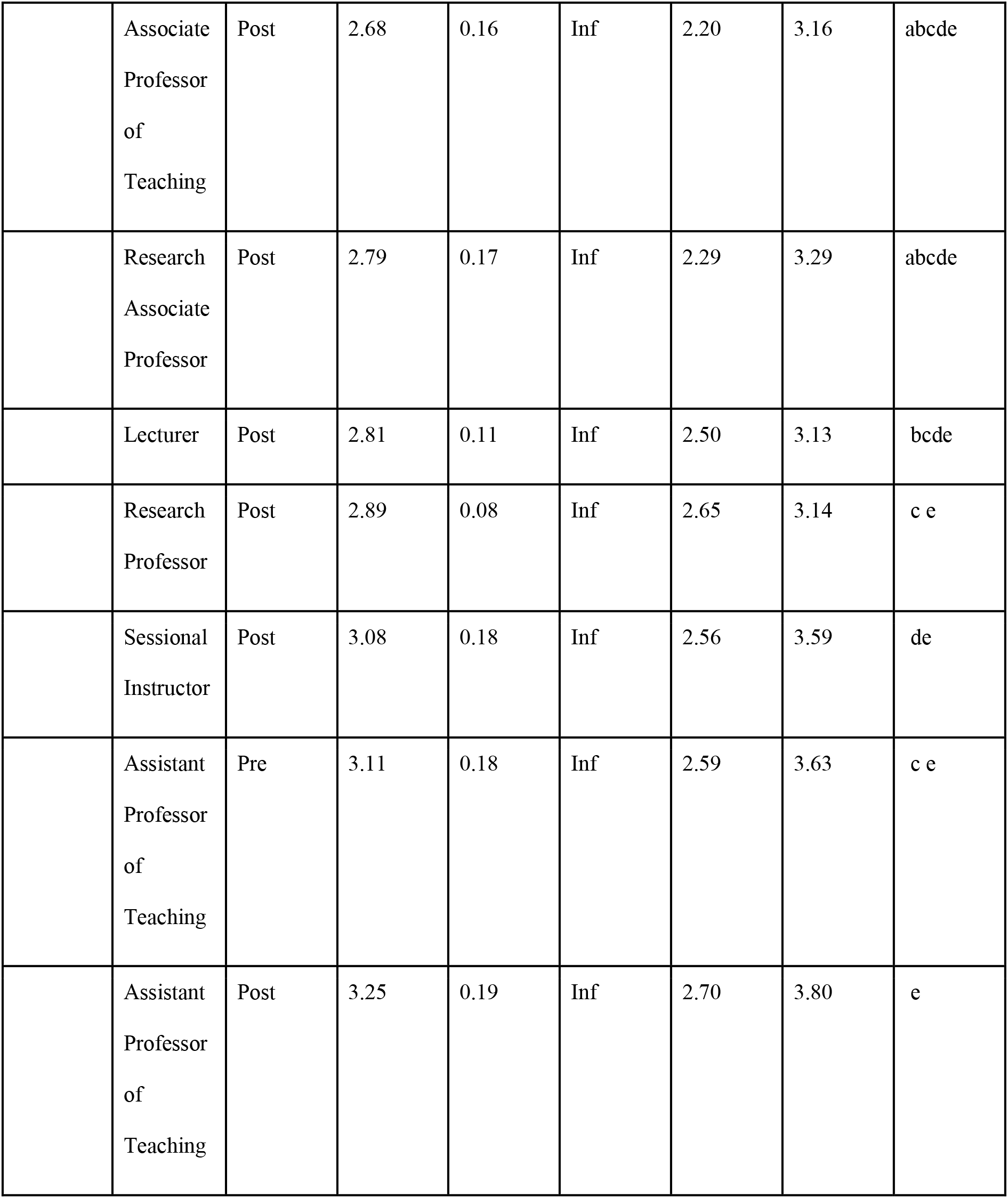
Post Hoc test results using sidak comparison, confidence level = 0.95, significance level used: alpha = 0.05. Letters indicate grouping.

**Table 3.** Tukey Post Hoc Test of only pre-term and post-term comparison. Bold values represent significant differences (p = 0.05).

| Pre-Post | Q1 | Q2 | F1 | F2 | F3 | F4 | F5 | F6 | F7 |
| --- | --- | --- | --- | --- | --- | --- | --- | --- | --- |
| Research Professor | **<.0001** | **<.0001** | **<.0001** | **<.0001** | **<.0001** | **<.0001** | **<.0001** | **<.0001** | **<.0001** |
| Associate Professor of Teaching | **<.0001** | 0.9914 | **0.0150** | 0.3147 | **0.0003** | **<.0001** | **<.0001** | 0.9732 | 0.9004 |
| Lecturer | **<.0001** | **<.0001** | **<.0001** | **<.0001** | **<.0001** | **<.0001** | **<.0001** | 0.8627 | **0.0013** |
| Postdoctoral Research Fellow | **<.0001** | **<.0001** | **0.0002** | 0.9826 | **0.0001** | **<.0001** | **0.0221** | 0.9985 | 0.9956 |
| Research Associate Professor | **<.0001** | **<.0001** | **0.0167** | **0.0215** | **0.0058** | **<.0001** | 0.3965 | 0.6989 | 0.9718 |
| Assistant Professor of Teaching | **<.0001** | **<.0001** | **0.0353** | 0.5276 | **0.0328** | **<.0001** | **0.0136** | 0.9028 | 1.0000 |
| Sessional<br>Instructor | **<.0001** | **<.0001** | **<.0001** | 0.9785 | 0.1201 | **0.0002** | **<.0001** | 0.6779 | **0.0001** |

